# Synergizing Network Pharmacology and Pan-Cancer Analysis in TCM Repositioning for Tumor Therapy

**DOI:** 10.1101/2025.01.03.631162

**Authors:** Dongli Yang, Xiyun Hu, Yong Liu, Yong Liao, Aamir Fahira, Lishan Wang, Muhammad Shahab, Defang Ouyang, Weihong Kuang, Zunnan Huang

## Abstract

**Background:** Repositioning Traditional Chinese Medicine (TCM) for cancer treatment, addressing heterogeneity through synergistic effects, aligns with the evolving trend of combination therapy. However, the complexity of TCM and the lack of methodology hinders the elucidation of TCM treatment mechanisms and potential indications. The research aims to construct develop a comprehensive method combining network pharmacology and pan-cancer analysis (NetPharm-PanCan) for TCM repositioning, exemplified by the Astragali Radix-Curcumae Rhizoma (ARCR) herb pair.

**Method:** The TCM-component-gene network was constructed using Cytoscape3.7.2 with gene screening based on components from TCMSP and targets predicted via the Swiss Target Prediction database. The core gene set of the ARCR (CGS_ARCR_) was identified by analyzing network via the dual algorithm, namely Degree and MCODE followed by GO and KEGG enrichment analyses. The pan-cancer analysis encompassing Gene Set Variation Analysis (GSVA), immune infiltration correlation, cancer pathway analysis, and Gene Set Enrichment Analysis (GSEA) was unveiled to identify the potential indications. Multivariate cox regression analysis was employed for identifying prognostic genes, followed by the modeling for potential therapeutic indications using the R software. The Metescape database was harnessed to reveal similarities and differences in the mechanisms for prognostic genes associated with potential indications. The XGBoost algorithm was used to construct an indication prediction system according to the analysis results above.

**Result:** The CGS_ARCR_ comprises 28 genes targeting all ARCR components. The pan-cancer analysis revealed that the CGSARCR showed significantly higher tumor scores than normal tissue scores across nine different types of cancer, including THCA, KIRC, LUAD, COAD, BRCA, STAD, ESCA, and others. The CG_SARCR_ correlated strongly with cancer-related pathways and the immune microenvironment. Prognostic models evaluate the potential of these indications as follows: THCA> KIRC> LUAD> COAD> BRCA> STAD> ESCA, with enrichment analysis suggesting KIRC and LUAD as the most potential indications of ARCR. The XGBoost-based system achieved high predictive accuracy (training AUC: 0.985, testing AUC: 0.96, training logloss: 0.21, testing logloss: 0.25).

**Conclusion:** The NetPharm-PanCan method provides a robust, network-based pan-cancer analysis framework for TCM repositioning in cancer research. It provides theoretical foundations and practical tools for TCM-based drug development, with potential applicability to broader drug repurposing efforts.

## 1. Background

Cancer, as a heterogeneous disease, persists as a prominent factor contributing to mortality on a global scale, with approximately 10 million patients succumbing to this ailment each year ^1^. Cancer complexity arises from vast genetic and molecular variations within and across tumor types, making it difficult for single-target treatments to achieve broad effectiveness. This challenge highlights the need for therapies that can simultaneously target multiple pathways. With ongoing advancements in technology and a more comprehensive understanding of human tumor disorders, there is greater potential to minimize cancer mortality via innovative therapeutic approaches ^2,3^. However, the discovery of a novel anti-cancer compound requires a considerable cost of approximately $648 million and a long duration of several years to more than a decade ^4^, resulting in a drug development timeline far slower than anticipated ^5,6^. Drug repurposing is a widely acknowledged strategy in the realm of drug development, which exploits potential mechanisms of action and targets genes of existing drugs in old therapeutic areas to explore new therapeutic indications ^7,8^. This strategy substantially curtails the cost and time associated with cancer drug development and helps to alleviate drug shortages ^9^. It aims to revolve around the optimization of existing anti-cancer drugs, providing a valuable avenue for drug repurposing in precision oncology^10^.

The application of drug repurposing in cancer treatment initially focused on the investigation of individual Western pharmaceutical compounds ^11^. In recent years, several studies have reported some successful instances of drug repurposing for single drugs. For instance, Olaparib and Capecitabine have been repurposed to treat breast cancer and neuroblastoma, respectively ^12,13^. However, due to the high degree of inter-tumor and intra-patient heterogeneity, the use of single drugs often faces challenges in achieving sufficient cytotoxic effects when treating different types of tumor cells ^14–16^, highlighting the need for more comprehensive therapeutic strategies. The network pharmacology of TCM has rapidly evolved thanks to its potential to facilitate drug repurposing and develop multi-targeted drugs ^17,18^. This Network-based approach seeks to explore multi-component Chinese herbal compounds to address the heterogeneity of tumors more comprehensively and offer new prospects for drug repurposing strategies ^18^. In addition, compared to single drugs, TCM is believed to exhibit synergistic effects in cancer treatment, which can effectively counteract the heterogeneity of tumor cells through multiple components, target genes, and pathways, thereby reducing the side effects of traditional treatments and playing a crucial role in inhibiting proliferation as well as immunotherapy of cancers ^19^. Consequently, TCM repurposing represents a promising avenue teeming, poised to unlock the full potential of TCM in tumor treatment, probably becoming one of the novel directions for future anti-cancer therapy.

Astragali Radix and Curcumae Rhizoma are two extensively researched TCMs that have shown significant activity in the field of cancer therapy ^20,21^. Astragali Radix is characterized by its immunomodulatory and anti-angiogenic properties ^20^, while Curcumae Rhizoma is renowned for its remarkable anti-inflammatory and antioxidant activities ^21^. Because of their distinct and complementary mechanisms of action, there has been notable interest in exploring their combination. The Astragali Radix-Curcumae Rhizoma (ARCR) exhibited remarkable anti-tumor effects in clinical settings for various cancers, including lung cancer ^22^, colorectal cancer ^23,24^, gastric cancer ^25^, and breast cancer ^26^. Furthermore, numerous network pharmacology studies on these cancers have indicated that the ARCR can synergistically enhance their anti-tumor properties by acting through multiple targets and pathways to inhibit cancer development and progression ^27–29^. However, the theories and mechanisms underlying the use of ARCR remain incompletely elucidated. Thus, the pursuit of its pharmacological repurposing holds the promise of unraveling this riddle, affording a profound comprehension of its untapped antitumor potential and shedding light on the therapeutic commonalities and distinctions that span multiple cancers.

Although computational methods for drug repurposing, such as feature-based, network-based, machine-learning-based methods, and more have been widely reported and employed to study the repurposing of single drugs ^30^, it is difficult for these methods to systematically cope with the complexity and uncertainty of the pharmacological effects of TCM. Currently, research on the repurposing of TCM in cancer treatment is relatively scarce, due to a lack of systematic methods for repurposing TCM formulations. The emergence of bioinformatics fields including network-based pharmacology analysis, whole genome sequencing (WGS), and pan-cancer research has provided novel opportunities for therapeutic repurposing ^31^. Their combination can allow for the integration of clinical data and intricate herbal knowledge to unravel the intricate web of multi-component and multi-target systems, thereby reducing costs, facilitating the comprehensive exploration of the repurposing of TCM in the field of cancer and illuminating new vistas for research in complex system of TCM.

In this study, we introduce the NetPharm-PanCan approach, a novel framework combining network pharmacology and pan-cancer analysis with whole-genome sequencing (WGS) data, specifically tailored for TCM repurposing. Existing computational repurposing methods struggle to systematically capture TCM complex pharmacological profile. Our NetPharm-PanCan framework, by integrating network pharmacology with pan-cancer and WGS data, uniquely enables a comprehensive analysis of multi-component TCM formulations, paving the way for more efficient repurposing strategies. Specifically, we utilized network pharmacology analysis with MCODE and Degree algorithms to identify core genes of ARCR (CGS_ARCR_). Employing the core gene set, the large-scale RNA-seq data from TCGA were used to perform pan-cancer analysis for screening potential cancer indications. Additionally, we constructed prognosis models using multivariate COX regression analysis, which not only assessed the predictive performance of prognosis models across diverse tumor types but also validated the reliability of the CGS_ARCR_ for exploring new indications. Finally, enrichment analyses of the CGS_ARCR_ and the prognostic gene sets for different indications were performed to illustrate the similarities and differences in the mechanisms of the ARCR for the treatment of different cancers.

In conclusion, NetPharm-PanCan establishes both theoretical and practical groundwork for the repurposing and advancement of TCM. This introduces innovative approaches to exploring the correlation between drugs and diseases, particularly within the realms of “same disease, different treatment” and “different diseases, same treatment”. By harnessing the multi-targeted and synergistic effects of TCM formulations, the NetPharm-PanCan approach illuminates a promising path for precision oncology, addressing cancer’s heterogeneity and paving the way for future studies on TCM repurposing.

## 2. Result

### 2.1. A brief overview of study design and outcomes

This study leverages the NetPharm-PanCan computational framework to systematically repurpose ARCR by identifying its potential applications across diverse cancer types, supporting the development of targeted, multi-component TCM therapies. As illustrated in **Figure 1**, NetPharm-PanCan integrates network pharmacology with pan-cancer analysis to identify new therapeutic indications for TCM while validating established ones. This rigorous approach, which incorporates prognostic modeling and enrichment analyses, demonstrates the potential of NetPharm-PanCan to advance research and development in TCM repurposing. The framework comprises five critical steps, detailed below.

**Figure 1.**
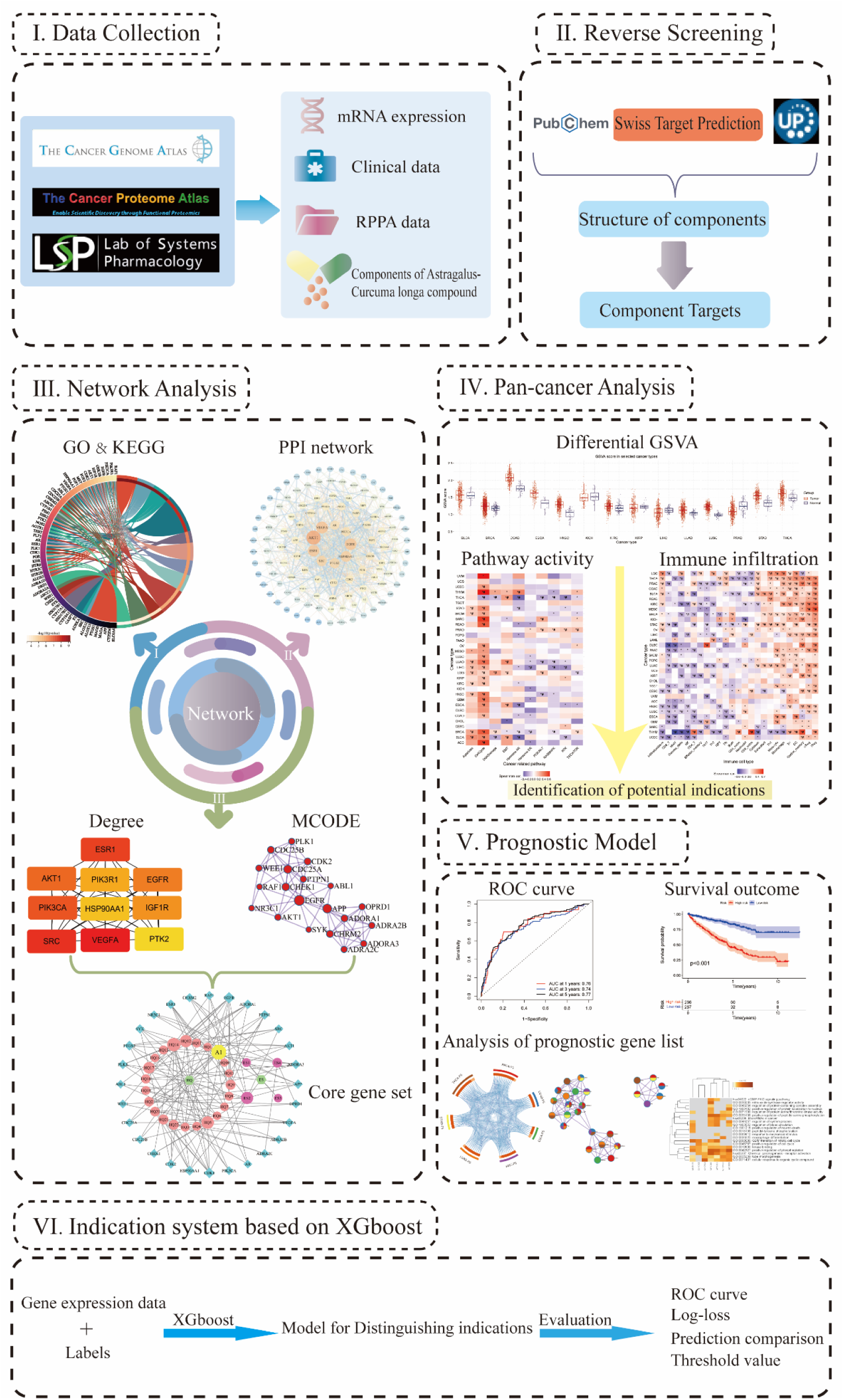
Schematic diagram of computational framework for the drug repurposing of ARCR.

The initial step involved extracting components of ARCR from drug-related literature and the Traditional Chinese Medicine Systems Pharmacology Database and Analysis Platform (TCMSP) database. In addition, The Cancer Genome Atlas (TCGA) database was used to obtain the gene expression data of pan-cancers along with clinical data (**Table S1**), whereas reverse phase protein array (RPPA) data was retrieved from The Cancer Proteome Atlas (TCPA) database. In the subsequent step, the targets related to these components were predicted through the reverse screening method. Moving on to step three, network pharmacology analysis was employed to construct a protein-protein interaction (PPI) network for the relevant target genes, followed by gene enrichment analysis, of which the results were further utilized to build a functional enrichment network of ARCR. MCODE and Degree algorithms were applied to analyze the functional enrichment network and PPI network, resulting in the core functional gene set and the core status gene set, separately. The combination of these two sets, with duplicate values removed, formed the core gene set of ARCR (CGS_ARCR_). In the most crucial fourth step, the CGS_ARCR_ and expression profile data were employed in comprehensive pan-cancer analysis to explicate the latent indications and mechanisms of ARCR at the pan-cancer level. Specifically, we conducted a Gene Set Variation Analysis (GSVA) followed by a Gene Set Enrichment Analysis (GSEA) to identify those cancers in which CGS_ARCR_ exhibited significant upregulation, thereby considering them as potential indications. Correlation analyses were performed to explore the associations of GSVA results with cancer pathways and immune microenvironments. Moving on to step five, prognostic models using clinical data for the potential indications were constructed to further evaluate their reliability. Enrichment analyses were conducted on various prognostic gene lists and CGS_ARCR_. These analyses aim to offer valuable insights into the underlying mechanism linked to potential indications relevant to cancer prognosis. Finally, we developed an indication prediction scoring system based on the XGBoost algorithm. This system can distinguish whether certain cancers are indications for ARCR, providing quantification and validation for the NetPharm-PanCan approach.

### 2.2. Active components and potential target genes of ARCR

To prepare for network pharmacology analysis, a total of 27 components of the ARCR were obtained. Specifically, 23 components were retrieved from the TCMSP database and an additional 5 components, namely Astragaloside I, Astragaloside II, Astragaloside III, Astragaloside IV, and Astragalus polysaccharides, were supplemented from the related literature ^26–29^. Among them, there are 23 active compounds of Astragali Radix (AR), including flavonoids, saponins, polysaccharides, amino acids, etc. while there are 5 active compounds of Curcumae Rhizoma (CR), including curcumin, flavonoids, volatile oils, polysaccharides, and hederagenin which is the common component (**Table 1**). Utilizing the Swiss Target Prediction repository, we conducted target prediction for the 27 active components, which yielded a total of 442 targets.

**Table 1.**
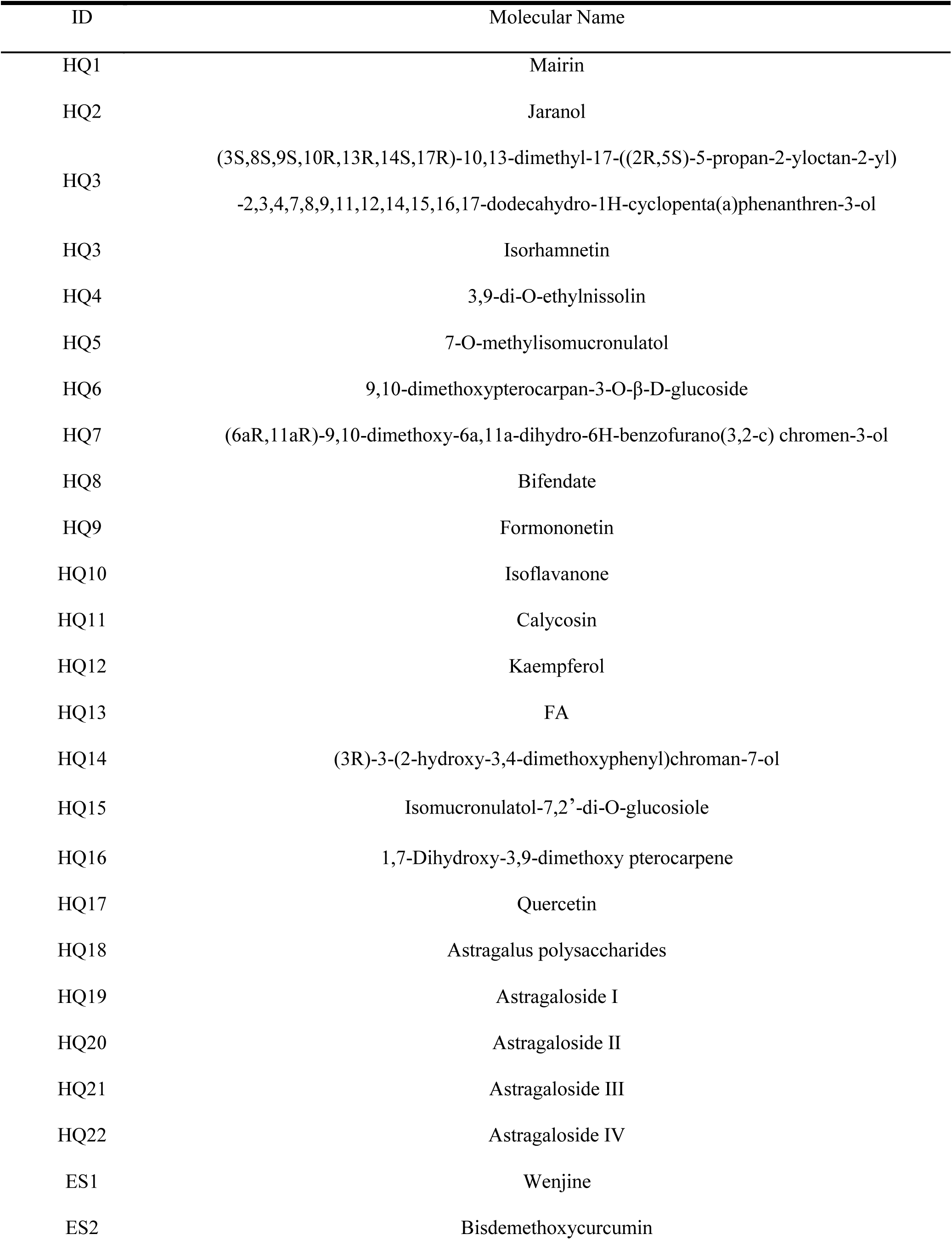
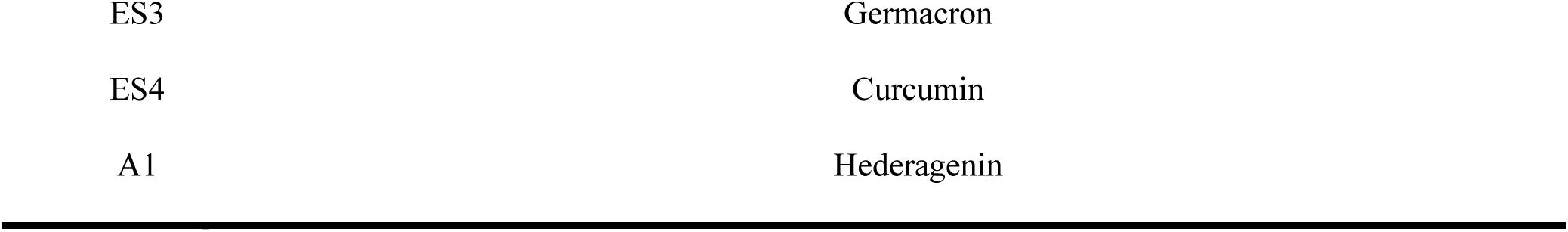
Active components and labels of Astragali Radix-Curcumae Rhizoma.

### 2.3. ARCR-component-target gene network and PPI networks

**Figure 2** shows the ARCR-component-target gene network constructed by Cytoscape version 3.7.2. The network structure highlights the complex relationships among herbs, components, and target genes. Notably, the shared component A1 (Hederagenin) has a high degree of connectivity, ranking fifth among all components, which signifies its potential importance in the network. Among the AR components, HQ5 (7-O-methylisomucronulatol), HQ9 (Formononetin), HQ12 (Kaempferol), HQ14 ((3R)-3-(2-hydroxy-3,4-dimethoxyphenyl) chroman-7-ol), and HQ17 (Quercetin) display the highest degree centrality, indicating their substantial influence on multiple target genes. In the CR section, ES2 (Bisdemethoxycurcumin) and ES4 (Curcumol) stand out as the most connected components, further emphasizing their relevance within the network. The remaining components likely play complementary roles, enhancing the therapeutic potential of the ARCR.

**Figure 2.**
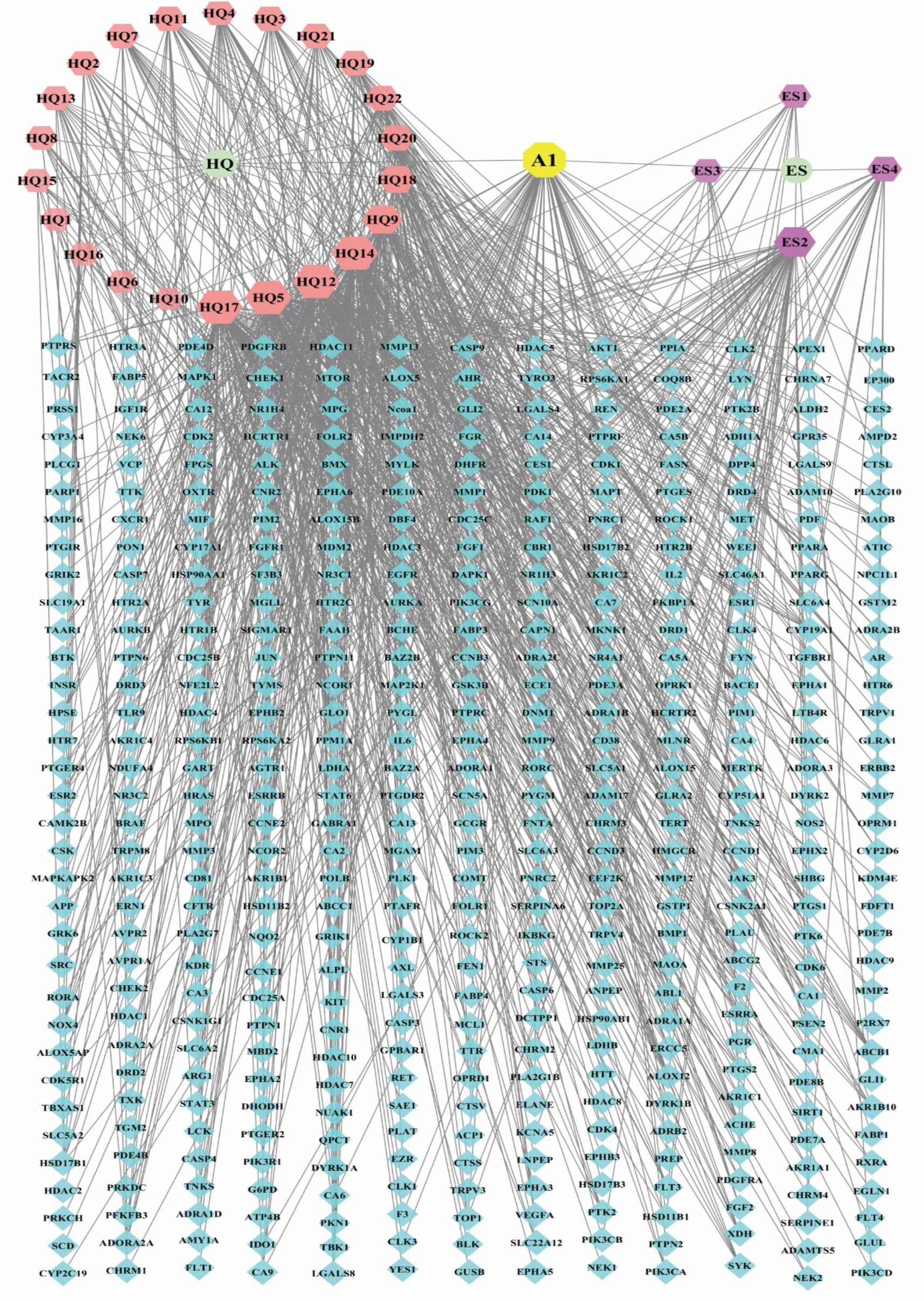
ARCR-component-target gene network. The network comprises a total of 471 nodes and 1036 edges. In the diagram, circles represent herbs (two in total), hexagons represent active components (27 in total), an octagon represents the shared component (one in total), and rhombuses represent component target genes (442 in total). The size of each hexagon reflects the degree of connectivity, with larger hexagons indicating components that are more centrally connected to multiple target genes, thereby underscoring their significance.

Building upon the ARCR-component-target gene network, **Figure 3** presents the PPI network of ARCR, focusing on the interplay among the most central target genes. From the initial pool of 442 target genes (**Table S2**), 125 target genes were selected utilizing the median degree value as the threshold within the ARCR-component-target gene network, with four genes being excluded due to a lack of associations with other target genes. Consequently, the final PPI network of ARCR comprised 121 target genes and 802 edges (**Figure 3 and Table S3**). Discerned from the PPI network, genes such as AKT1, VEGFA, EGFR, ESR1, HSP90AA1, PTGS2, SRC, PIK3CA, and AR(gene) may serve as critical hub genes mediating various cancers through the ARCR.

**Figure 3.**
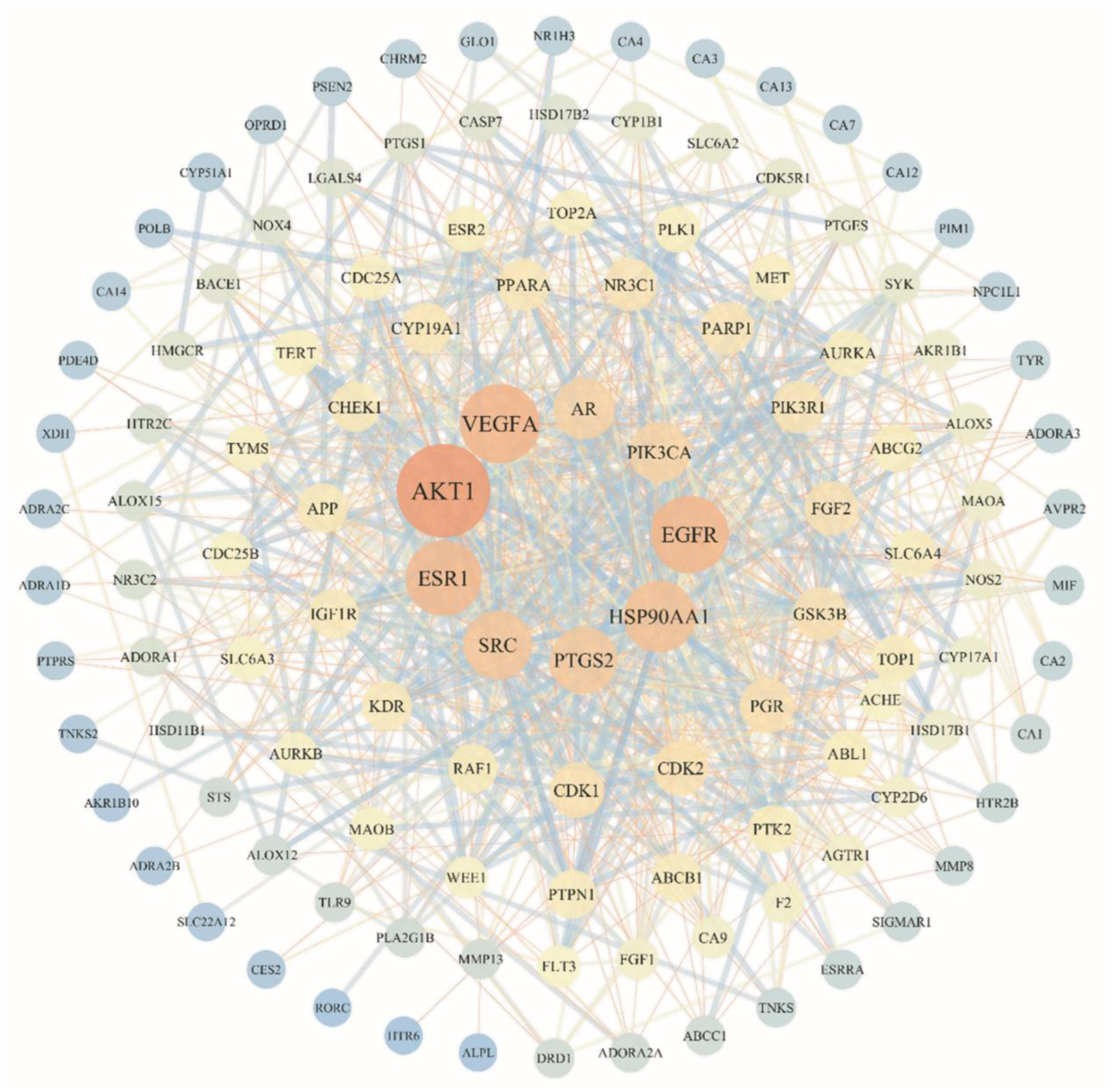
PPI network of ARCR. This PPI network represents differential centrality based on degree values. Brighter colors and larger node sizes indicate higher centrality, marking them as significant hubs in the network.

### 2.4. Performing Gene Ontology (GO) and KEGG Enrichment Analysis on PPI Network

A total of 1235 GO Biological Process (BP) terms, 94 GO Cellular Component (CC) terms, and 145 GO Molecular Function (MF) terms were identified by GO enrichment analysis on the 121 genes derived from the PPI network. All of these terms displayed significance levels with *P* < 0.05 (**Supplementary Sheet 1-3**). **Figure 4** depicts the top ten significant GO terms for BP, CC, and MF. Among these, the top ten GO-BP terms are predominantly associated with biological processes such as signaling, metabolism, cellular responses, and enzyme regulation. Notably, there is a tendency toward positive regulation of biological processes, including positive regulation of kinases, phosphorylation, and MAPK cascades (**Figure 4A**). The top ten GO-CC terms are associated with the organization of cell membranes and neuronal structures, suggesting potential influences of the PPI network genes on cell membrane composition and the nervous system (**Figure 4B**). As for the top ten GO-MF terms, they encompassed a wide range of enzyme activities, including protein kinase and phosphotransferase activities, oxidoreductase activities, and hydrolase activities, all of which play critical roles in biological processes (**Figure 4C**).

**Figure 4.**
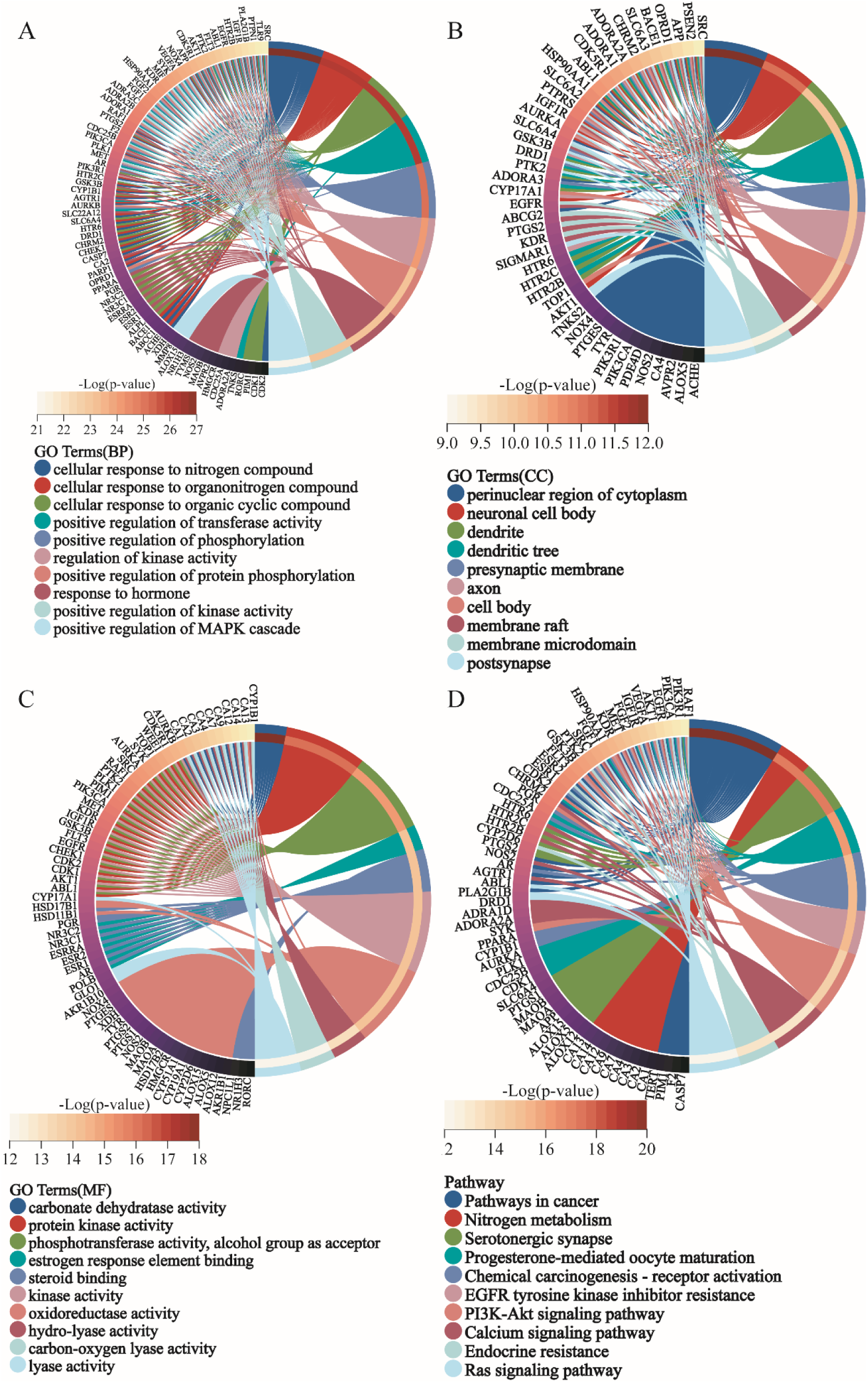
GO terms and KEGG pathways sorted by *P*-value in the PPI network of ARCR. Circle plots of the top ten **(A)** GO Biological Process. **(B)** GO Cellular Component, **(C)** GO Molecular Function, and **(D)** KEGG Pathway.

Furthermore, 121 genes from the PPI network were analyzed using KEGG pathways, which led to the identification of 146 pathways that were significantly enriched (*P* < 0.05) (**Supplementary Sheet 4**). Concisely, Pathways in cancer, Nitrogen metabolism, Serotonergic synapse, Progesterone-mediated oocyte maturation, Chemical carcinogenesis-receptor activation, EGFR tyrosine kinase inhibitor resistance, PI3K-Akt signaling pathway, and others were among the top 10 enriched KEGG pathways (**Figure 4D**). These pathways encompass a spectrum of processes, which are pertinent to cancer, metabolism, neurotransmission, reproduction, and cellular signaling, highlighting their multifaceted implications.

### 2.5. Core gene set of ARCR

A functional enrichment network is created by analyzing the enrichment of 121 target genes derived from the PPI network. This network assigns distinct biological significance to the input genes. Notably, only 66 genes demonstrate statistically significant biological relevance in this context (**Figure. 5A**). Eight distinct subgroups based on different functional categories are clustered from the 66 target genes applying the MCODE algorithm (**Figure 5B and Table S4**). Among these subgroups, MCODE1 emerges prominently, featuring the highest number of clustered genes and the top overall score within the 66-target genes network (**Table S4**). Remarkably, its top three significantly enriched KEGG pathways include the Cell cycle, cGMP-PKG signaling pathway, and Progesterone-mediated oocyte maturation (**Figure 6A and Table S5**). These pathways are frequently associated with cancer signaling pathways, particularly the Cell cycle which plays a crucial role in the development of various types of cancers. This association underscores the potential of this subgroup to intervene in cancer progression and makes it more likely to have therapeutic implications for multiple cancer types of ARCR. Therefore, the 20 genes of MCODE1 constituted the core functional gene set of the ARCR.

**Figure 5.**
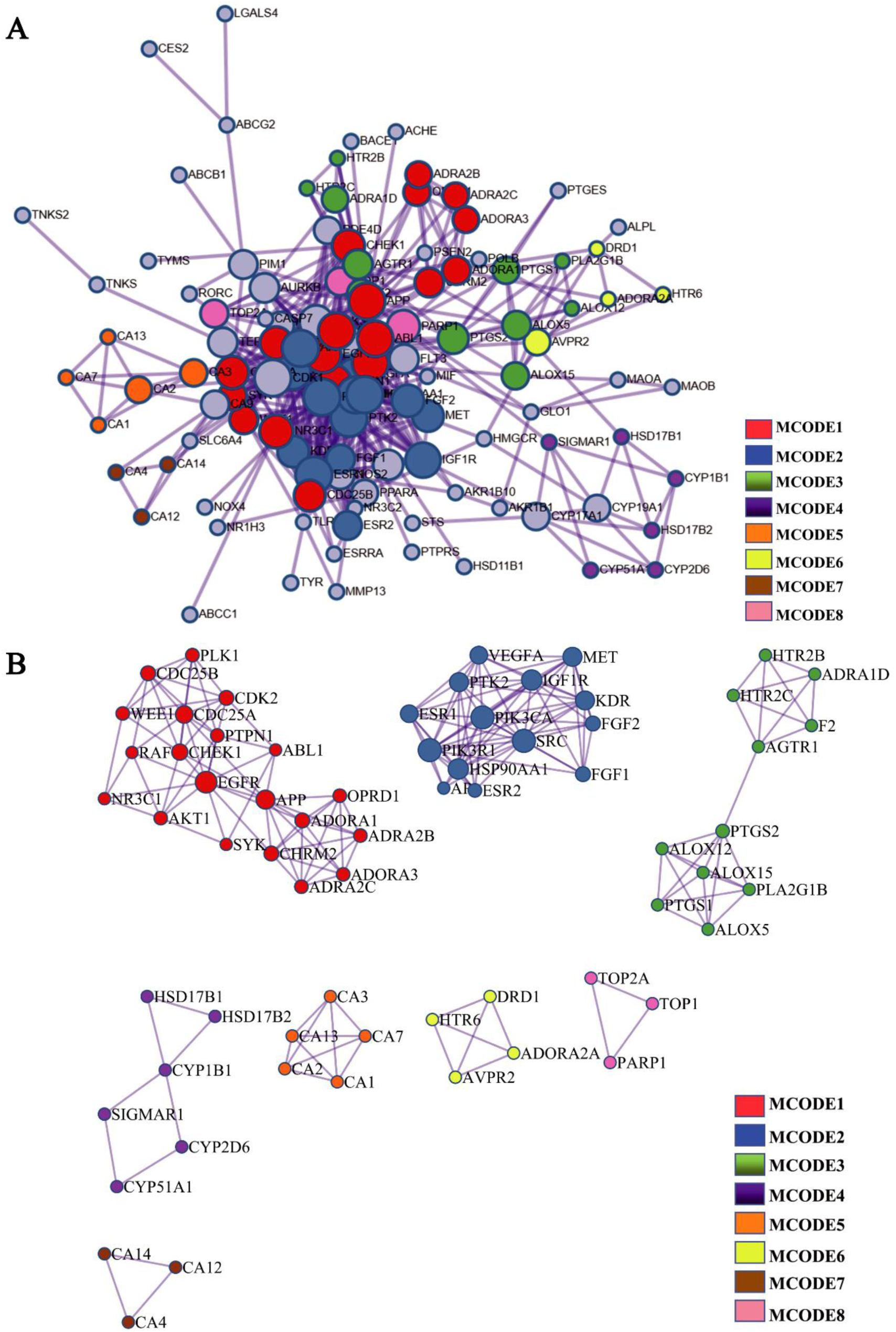
Analysis of the functional enrichment network of ARCR. **(A)** Protein functional enrichment network. **(B)** Eight protein functional enrichment subgroups obtained using the MCODE algorithm based on K-means clustering.

**Figure 6.**
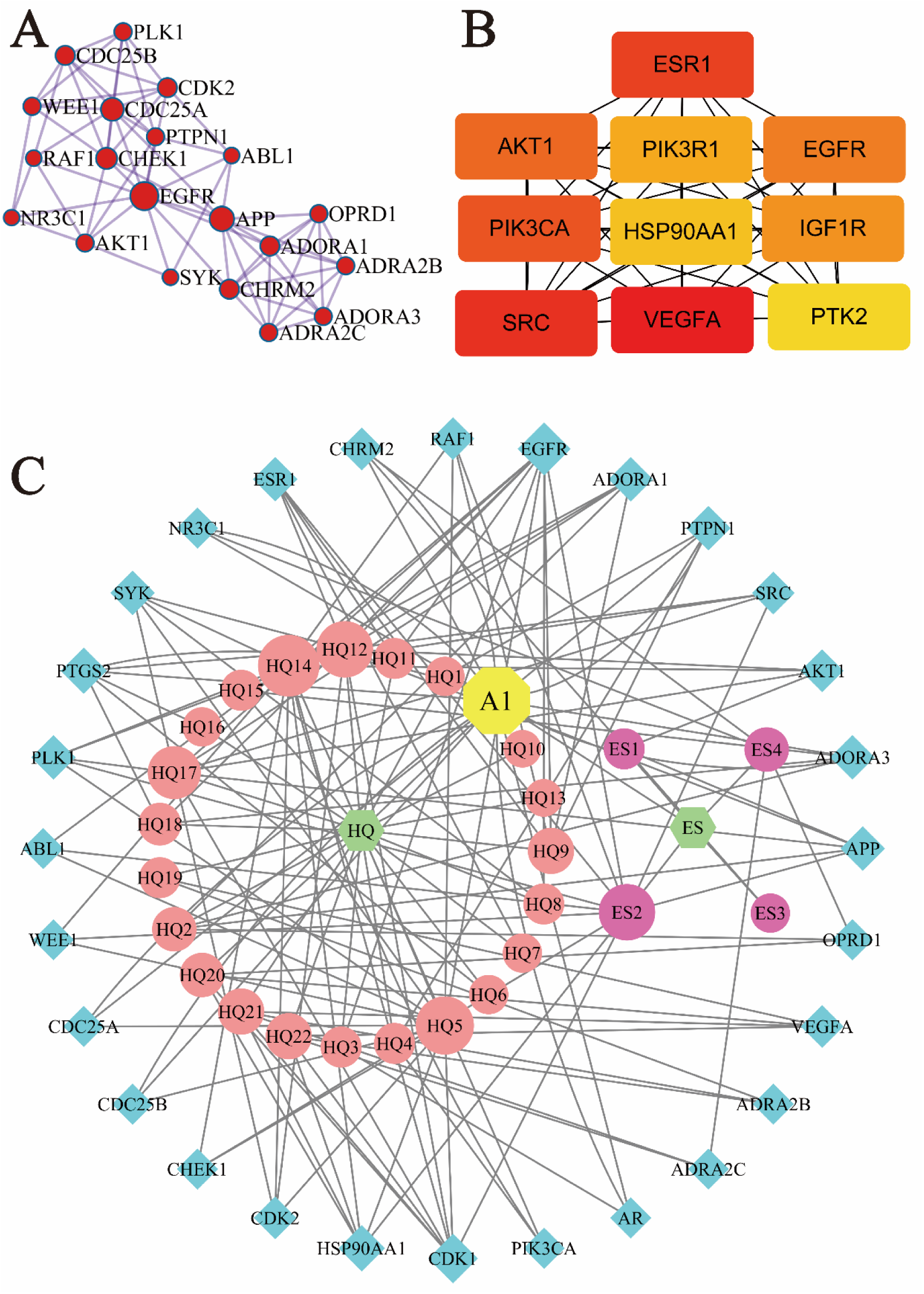
Construction of the core gene set for the ARCR(CGS_ARCR_). **(A)** The core functional gene set with the highest score (MCODE1) based on the MCODE algorithm in the core protein functional network. **(B)** Core status gene set of the ARCR, obtained by analyzing the PPI network using the Degree algorithm. **(C)** Herb-target gene network of the CGS_ACAR_ In the diagram, 28 blue diamonds represent the members of the CGS_ARCR_, 25 circles represent active components, one yellow octagon represents shared active components between AR and CR, and two green hexagons represent herbs.

The top ten target genes with the highest degree values are identified by analyzing the PPI network through the implementation of the Degree algorithm (**Figure 6B**). Higher degree values indicate a more prominent position within the PPI network. Thus, these genes constitute the core hub gene set of ARCR. Combining the core functional genes with the core status genes, the core gene set of ARCR (CGS_ARCR_) is derived, which contains 28 genes after the exclusion of two duplicates, namely AKT1 and EGFR (**Table S6**). The 27 components of ARCR are closely linked to the CGS_ARCR_, with each component targeting at least one core gene (**Figure 6C and Table S7**). This intricate linkage underscores both the representative and biological functionality of the CGS_ARCR_.

### 2.6. Differences in GSVA scores among normal and cancer samples

The comprehensive analysis performed using Gene Set Variation Analysis (GSVA) across 14 different cancer types reveals nuanced expression differences between tumor and normal samples. Notably, except for Bladder Cancer (BLCA) and Kidney Chromophobe (KICH), where differences were not statistically significant, the expression changes in the remaining 12 cancer types were statistically significant (*, *P* < 0.05, **Figure 7A**). Particularly in cancers such as Esophageal carcinoma (ESCA), Head and Neck Squamous Cell Carcinoma (HNSC), Colon Adenocarcinoma (COAD), Lung Squamous Cell Carcinoma (LUSC), Stomach Adenocarcinoma (STAD), Thyroid Carcinoma (THCA), Lung Adenocarcinoma (LUAD), Kidney Renal Clear Cell Carcinoma (KIRC), and Breast Invasive Carcinoma (BRCA), tumor tissues exhibited markedly higher GSVA scores than their normal counterparts, visually differentiated by red and blue box plots for tumor and normal samples respectively in **Figure 7A**. This pattern underscores the potential role of CGS_ARCR_ in promoting oncogenesis through overexpression mechanisms, potentially influencing pivotal cellular processes involved in tumorigenesis, as further detailed in **Table S8**.

**Figure 7.**
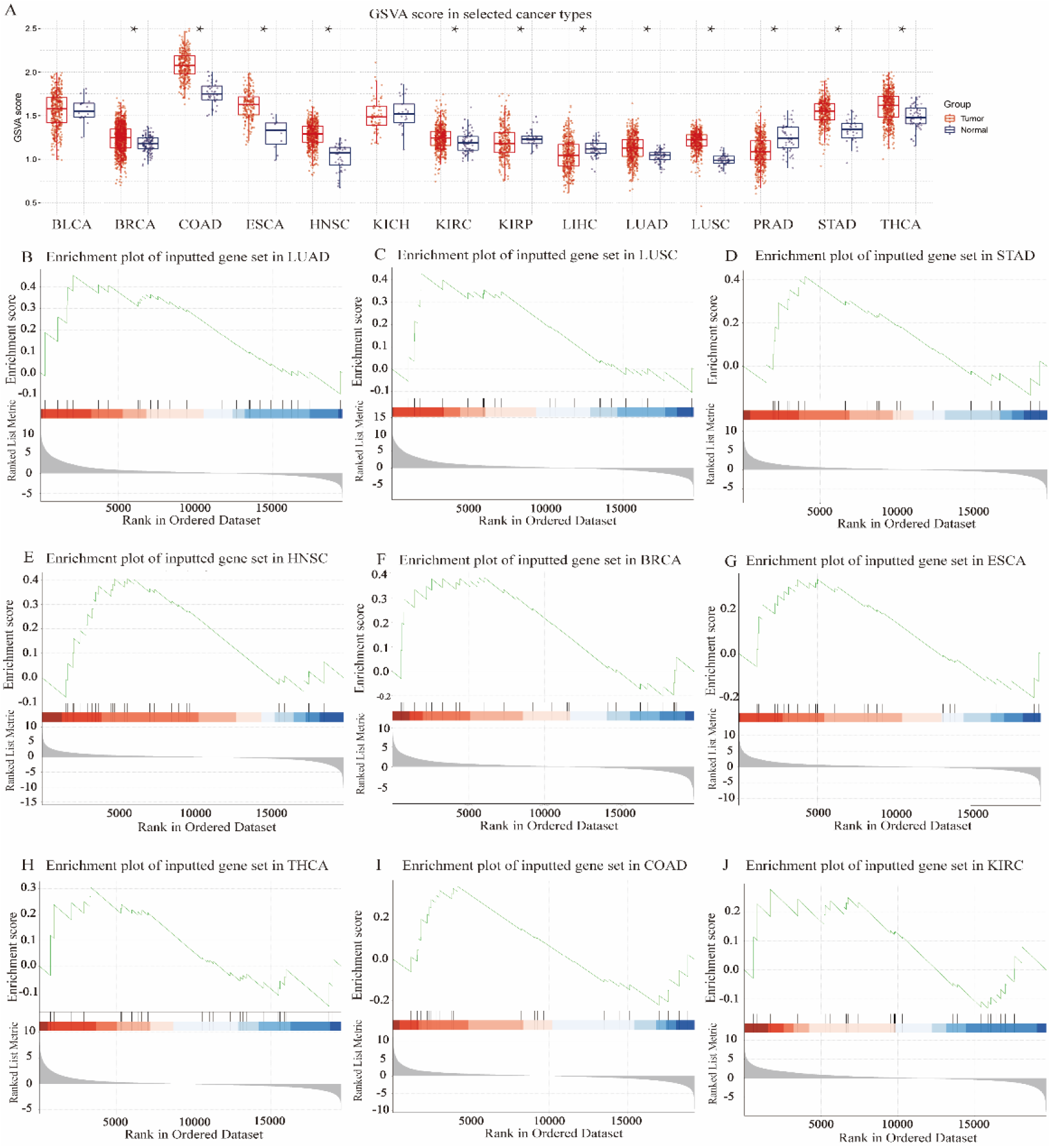
Differential Gene Set Variation Analysis and Gene Set Enrichment Analysis of CGS_ARCR_ Across Various Cancer Types. (**A**) GSVA scores presented as box plots to compare expression levels between tumor (red) and normal (blue) tissues across 14 types of cancer. Significant differences are denoted by asterisks (\**P* < 0.05), highlighting notable overexpression in tumor samples relative to normal samples in 12 out of 14 cancer types, except BLCA and KICH. (**B-J**) Gene Set Enrichment Analysis plots for selected cancer types (LUAD, LUSC, STAD, HNSC, BRCA, ESCA, THCA, COAD, and KIRC), demonstrating the correlation between CGSARCR gene set expression and cancer phenotypes. The top section of each plot shows the Running Enrichment Score (ES) with peaks indicating significant gene set correlations. The middle section maps the distribution of gene set members across a ranked list of genes (red bars indicate gene positions), and the bottom section depicts the ranking metric, transitioning from positive to negative values, which measures the correlation of gene expression with cancer phenotypes. These visual data corroborate the gene sets potential role in tumorigenesis and its implications for targeted cancer therapy.

### 2.7. GSEA at pan-cancer level

The gene set enrichment analysis (GSEA) of the CGS_ARCR_ revealed the distribution of enrichment scores (ES) in tumor tissues across nine potential indications (**Figure 7B-J)**. The ES for all nine potential indications is greater than 0, with the analysis ranking them in descending order as LUAD, LUSC, STAD, HNSC, BRCA, ESCA, THCA, COAD, and KIRC **(Table S9**). This indicates an overexpression of the CGS_ARCR_ in these cancers, which aligns well with the overexpression observed in GSVA by illustrating significant associations between CGS_ARCR_ and specific cancer phenotypes.

### 2.8. Correlation between GSVA Scores and Cancer Pathways

As shown in **Figure 8 (A)**, the GSVA scores of the CGS_ARCR_ exhibited significant positive correlations with various cancer pathways, including cell cycle, apoptosis, cell death, and epithelial-mesenchymal transition (EMT), suggesting that upregulating expression of the CGS_ARCR_ may activate these pathways in the specific cancer. Specifically, cell cycle was positively correlated with GSVA scores of all nine potential indications. Conversely, GSVA scores of the CGS_ARCR_ demonstrated significant negative correlations with estrogen receptor (ER) and RAS-MAPK pathways in some cancers, indicating a potential inhibitory effect of CGS_ARCR_ upregulation on these pathways. These findings underscore the significant correlations of the CGS_ARCR_ with key cancer pathways, suggesting its potential as the therapeutic target set by modulating specific pathways across various cancer types.

**Figure 8.**
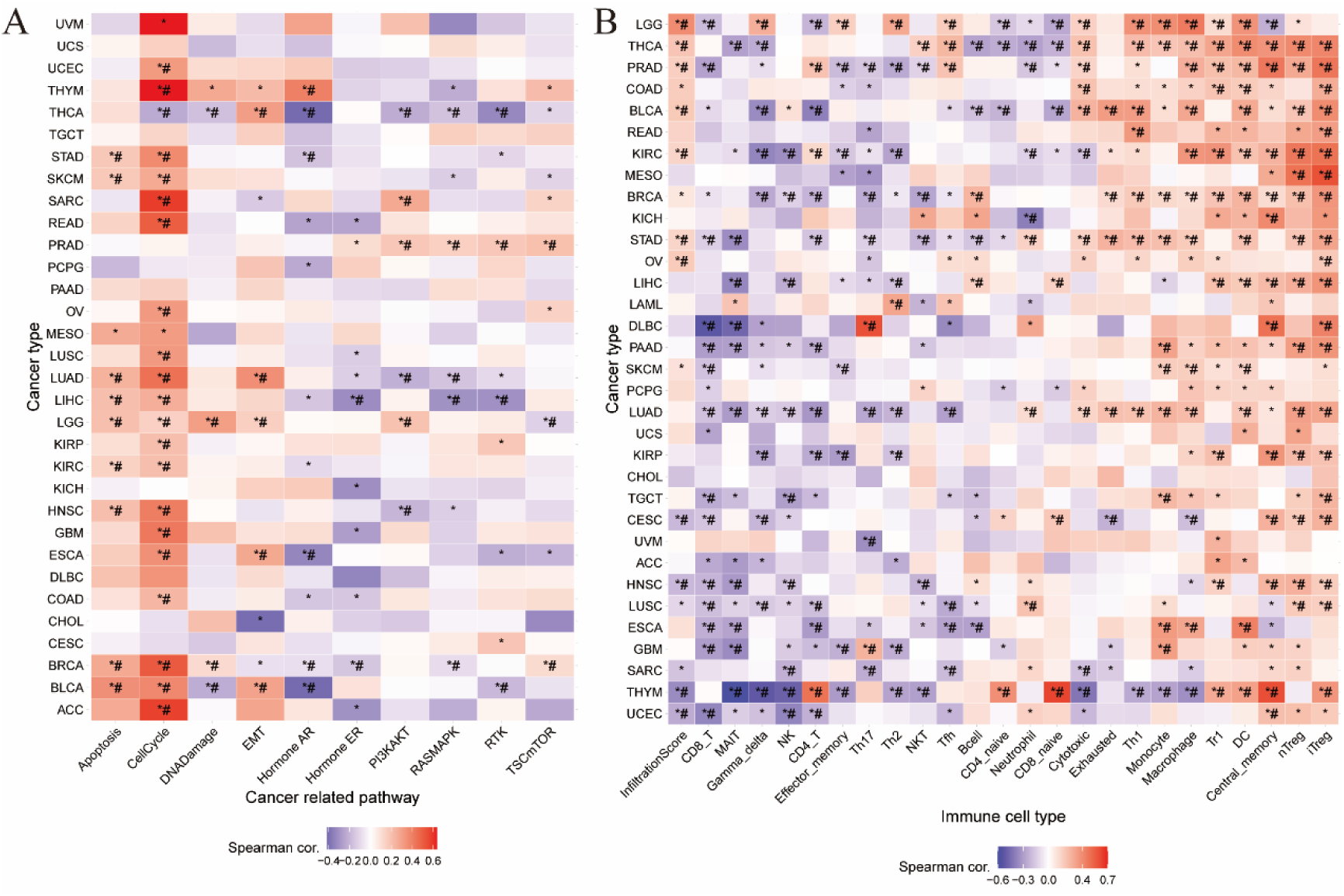
Correlation Analysis of the CGS_ARCR_. **(A)** Heatmap depicting the correlation between the GSVA scores of the CGS_ARCR_ and cancer-associated activity pathways in various cancers. **(B)** Heatmap illustrating the correlation between GSVA scores of the CGS_ARCR_ and immune cells in the microenvironment of different tumors. The sign * represents *P* value ≤ 0.05; # represents FDR ≤ 0.05. Red in the heatmap indicates a positive correlation, blue indicates a negative correlation, and the intensity of the color reflects the strength of the association between gene set expression and pathway correlation

### 2.9. Correlation between GSVA Scores and the Immune Microenvironment

The GSVA scores of the CGS_ARCR_ were positively correlated with various immune cells such as Tr1, DC, Central memory, nTreg, and iTreg in most cancers, while these scores showed a negative correlation with immune cells like CD8_T, MAIT, Gamma delta, NK, and CD4_T (**Figure 8B**). This indicated that the upregulation of the CGS_ARCR_ primarily enhances immune responses involving Tr1, DC, Central memory, nTreg, and iTreg while suppressing immune responses related to CD8_T, MAIT, Gamma delta, NK, and CD4_T. Such alterations suggest that CGS_ARCR_ plays a complex regulatory role in modulating immune functions, potentially influencing the immune infiltration processes within the tumor microenvironment.

### 2.10. Prognostic modeling of CGS_ARCR_ in potential indications

The prognostic models are constructed for nine potential indications using multivariate Cox regression analysis based on CGS_ARCR_. The number of prognostic genes in different cancers is as follows: THCA (13), BRCA (12), LUAD (10), KRIC (9), COAD (7), STAD (7), HNSC (7), ESCA (6), and LUSC (6) (**Table S10-S18**). **Table S10-S18** present the prognostic genes based on the nine cancers, along with their *P-values*, coefficients (coef), Hazard Ratios (HR), and 95% Confidence Intervals (CI).

**Figure 9** depicts the ROC curves for the prognostic risk models. Among these, the model of THCA stands out as the most reliable, followed by KIRC and LUAD, with their time-dependent AUC values for 1-7 years of survival exceeding 0.7. The AUC values for 1, 3, and 5 years are as follows: THCA (0.89, 0.91, 0.89), KIRC (0.76, 0.74, 0.77), and LUAD (0.72, 0.73, 0.72) (**Figure 9A-C**). These models can confidently and effectively predict adverse survival outcomes for corresponding cancer patients over an extended period. The models constructed for COAD, BRCA, ESCA, and STAD showed AUC values exceeding 0.7 for three consecutive years, demonstrating strong predictive performance (**Figure 9D-G**). These prognostic models can confidently predict adverse survival outcomes for COAD patients for 4-7 years, BRCA patients for 3-5 and 7 years, ESCA patients for 3-5 years, and STAD patients for 4-6 years. This suggests that the models predict outcomes precisely across the mentioned time spans. In contrast, LUSC and HNSC exhibited AUC values below 0.7 for 1-7 years, indicating moderate predictive performance (**Figure 9H, I**). Their models have limitations in predicting clinical outcomes for patients. Therefore, these two types of cancer are excluded from the potential indications.

**Figure 9.**
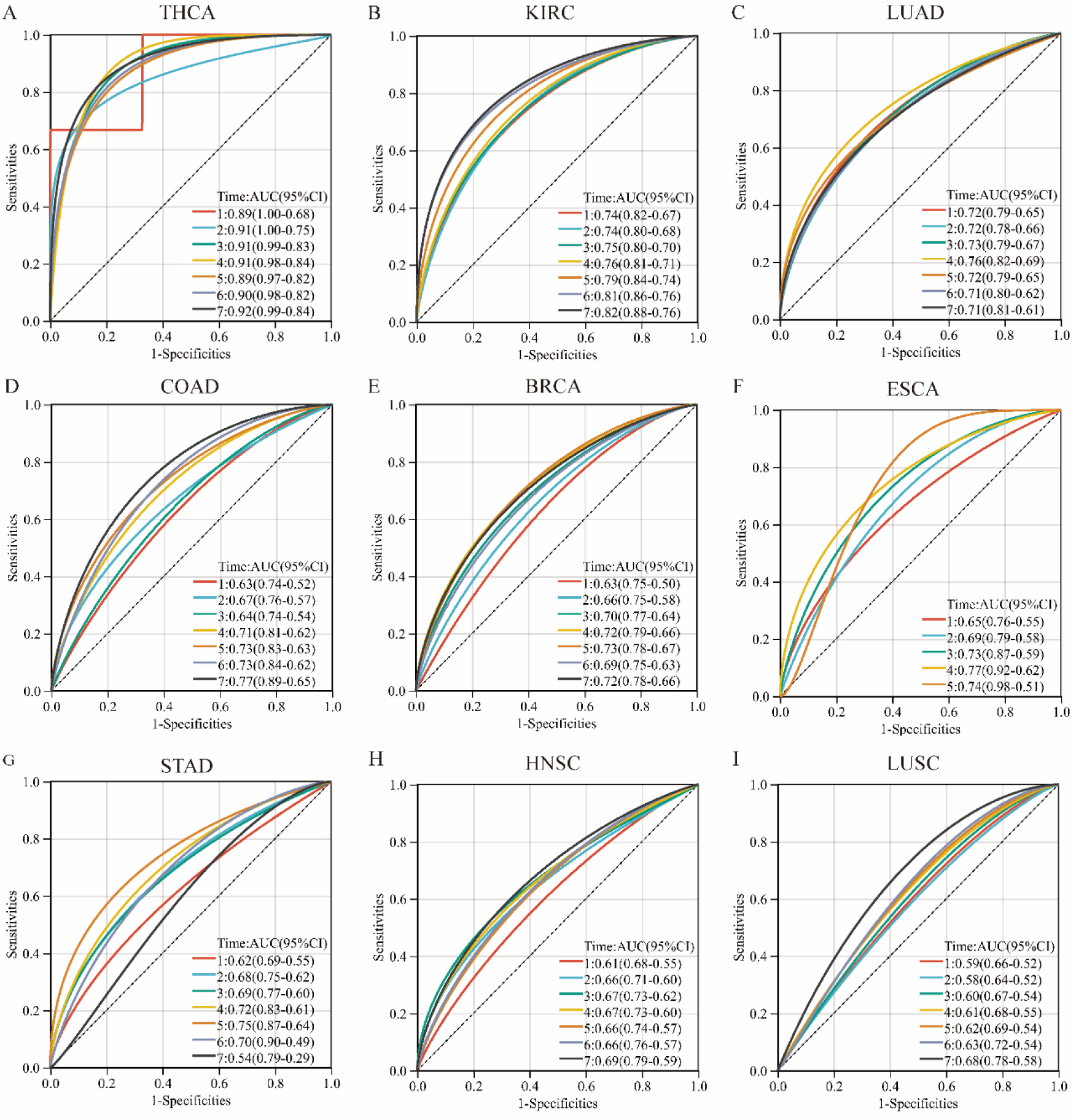
Prognostic Models of the CGS_ARCR_ in Nine Potential Cancers. ROC curves of the Prognostic model for the CGS_ARCR_ in **(A)** THCA, **(B)** KRIC, **(C)** LUAD, **(D)** COAD, **(E)** BRCA, **(F)** ESCA, **(G)** STAD, **(H)** HNSC, and **(I)** LUSC.

**Figure 10 (A-I)** presents the survival analysis curves for the prognostic models of the nine cancers. These Kaplan-Meier survival curves consistently show that the low-risk group has significantly higher survival rates than the high-risk group, with all *P*-values less than 0.05. These findings indicate that the prognostic models constructed based on CGS_ARCR_ can effectively assess the prognosis risk of cancer patients. They exhibit high accuracy in predicting the survival outcomes of cancer patients, and the results of the Kaplan-Meier survival curves further validate the efficacy of the prognostic models.

**Figure 10.**
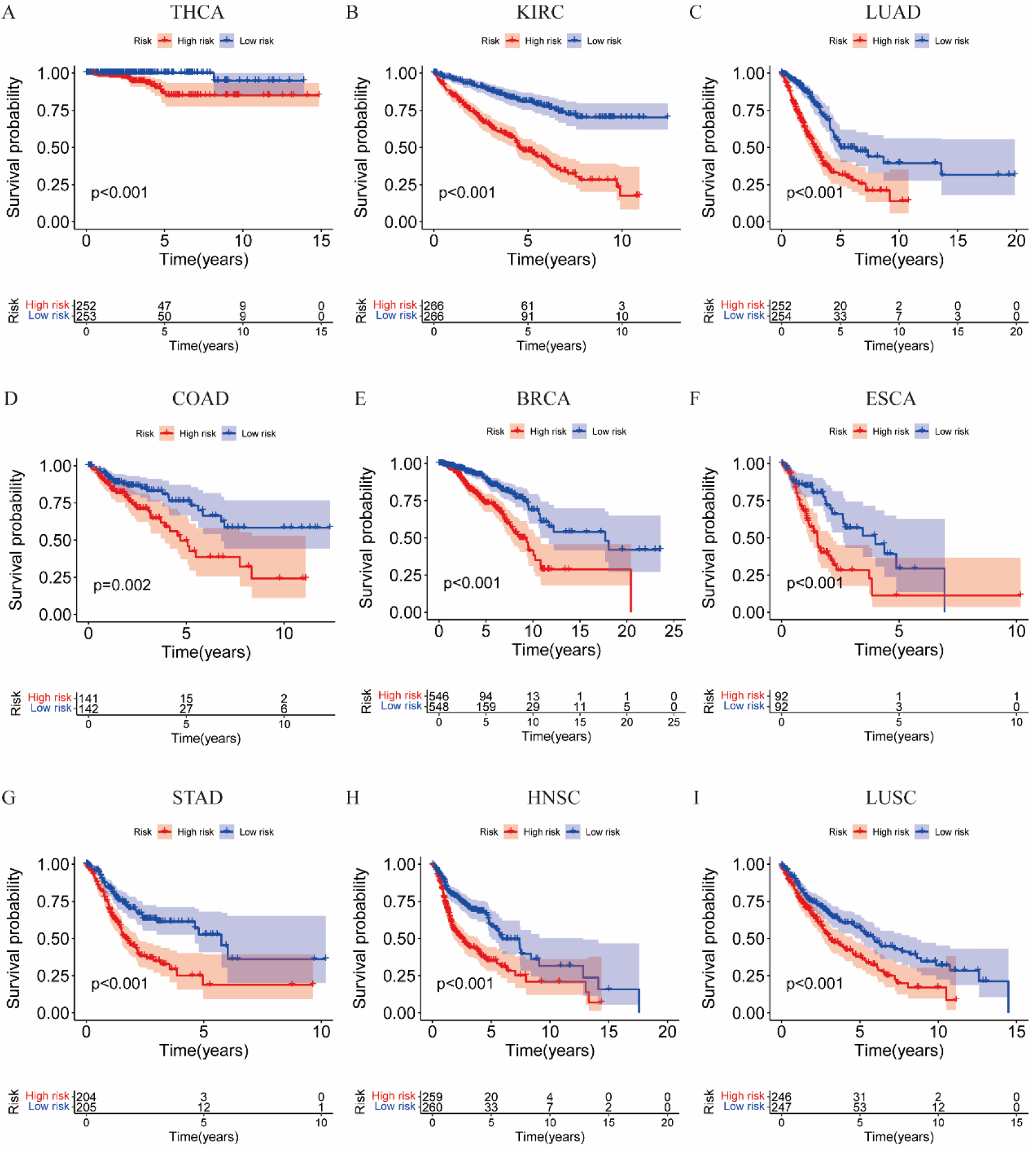
Survival curves of CGS_ARCR_ in potential prognostic models for nine different cancers. K-M survival curves in **(A)** THCA, **(B)** KRIC, **(C)** LUAD, **(D)** COAD, **(E)** BRCA, **(F)** ESCA, **(G)** STAD, **(H)** HNSC, and **(I)** LUSC.

### 2.11. Similarities and differences in prognostic genes for seven reliable potential indications

**Figure 11** illustrates the similarities and differences among the potential indications with reliable prognoses associated with the ARCR. The circos plot of gene association displayed an overlap among most of the prognostic genes from the seven potential cancer indications, indicating a genetic-level similarity in the mediated effects of ARCR on these seven cancers (**Figure 11A**). In addition, the other circos plot of pathway association derived from the gene associations, demonstrated a high functional pathway similarity of ARCR in these seven cancers among the prognostic genes (**Figure 11B**). Furthermore, the PPI network of the prognostic genes of the seven cancers revealed the presence of several shared genes across the prognostic gene lists (**Table S19**). These genes could simultaneously influence the prognoses of multiple cancers, including ADRA2C (THCA, KIRC, LUAD, BRCA, STAD); ESR1 (THCA, KIRC, COAD, STAD, BRCA); HSP90AA1 (THCA, KIRC, BRCA, ESCA); PLK1 (KIRC, LUAD, COAD, BRCA); CDC25A (THCA, BRCA, ESCA); CDC25B (THCA, LUAD, ESCA); NR3C1 (LUAD, COAD, STAD); AR (KRIC, STAD), and more (**Figure 11C**). Notably, ADRA2C, ESR1, HSP90AA1, and PLK1 appeared in the prognostic lists of four or more cancers, suggesting their widespread involvement in the pathways targeted by ARCR in the therapy of these seven cancers.

**Figure 11.**
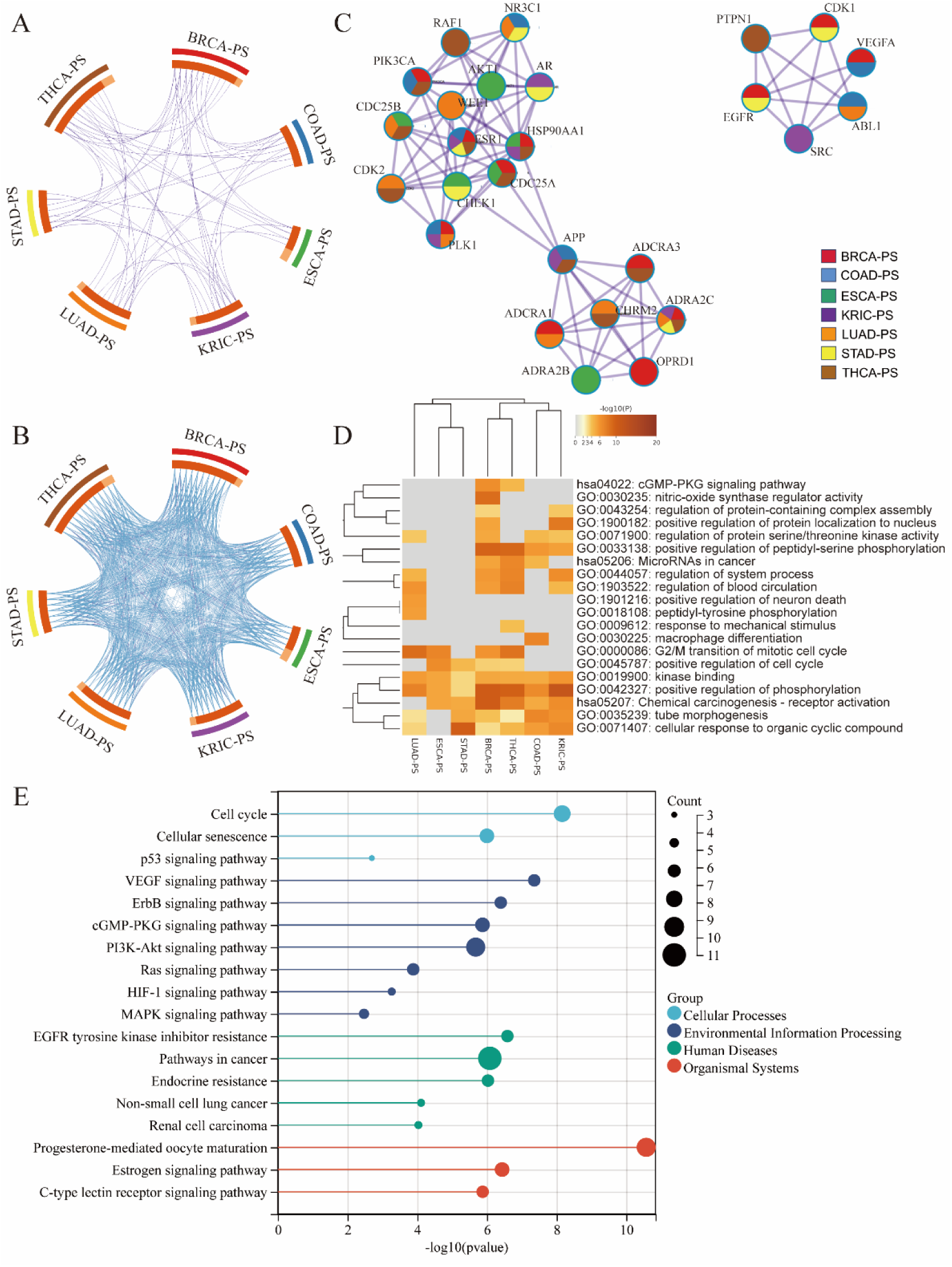
Comparative Analysis of Seven Potential Cancer Prognostic Gene Lists Targeted by the ARCR. **(A)** Circos plot illustrates gene overlap among the seven gene lists. Outer arcs in different colors represent the identity of different gene lists, while inner arcs represent the gene lists, with each gene member assigned a position on the arc. Deep orange indicates genes shared across multiple lists, and light orange represents genes unique to that particular list. Purple lines connect common prognostic genes among different cancers at the gene level. **(B)** Building upon (A), blue lines signify genes in different prognostic gene lists sharing common pathways. **(C)** PPI network of prognostic gene lists for the seven cancers, displaying the distribution of their common genes. **(D)** Comparative analysis of GO and KEGG terms among prognostic gene lists of the seven cancers. **(E)** KEGG enrichment analysis of the CGS_ARCR_

**Figure 11D** presents the differences and similarities among prognostic gene lists from seven cancer indications analyzed through GO and KEGG enrichment. Despite the absence of genes that simultaneously served as prognostic markers for the seven prognostic genes list, their gene compositions exhibited a high degree of similarity in function, resulting in significant consistency in biological processes such as kinase binding and positive regulation of phosphorylation (**Figure 11D)**. Among these processes, the most prominent participation is observed in BRCA and THCA. Concerning KEGG pathways, except for the prognostic genes list of LUAD, the prognostic lists of the other six cancer types were all involved in Chemical carcinogenesis-receptor activation. ESR1 is a shared gene among KRIC, COAD, STAD, BRCA, and THCA (**Table S19**). Thus, these findings suggest that ESR1 may play a crucial role in chemical carcinogenesis through receptor activation, potentially linked to estrogen receptor α activation. Moreover, the prognostic gene lists of BRCA, COAD, and THCA were associated with microRNAs in cancer. Besides these shared components, some cancer types also exhibited significantly enriched terms respectively. For instance, LUAD-PS (the prognosis genes of LUAD) was involved in positive regulation of neuron death, BRCA-PS was associated with nitric-oxide synthase regulator activity, THCA-PS was connected to response to mechanical stimulus, and COAD-PS was networked with “macrophage differentiation (**Figure 11D**). These distinct enrichment processes may stem from the differences in prognostic gene lists, further highlighting the relationship between the distinctiveness of prognostic gene lists and biological function.

**Figure 11(E)** depicts the KEGG enrichment analysis of the CGS_ARCR_. The results demonstrated that the CGS_ARCR_ primarily enriched four major categories, comprising Cellular Processes, Environmental Information Processing, Human Diseases, and Organismal Systems. The significant pathways, each containing the highest number of genes, were Pathways in cancer, PI3K-AKT signaling pathway, Progesterone-mediated oocyte maturation, and Cell cycle. Additionally, in terms of human diseases, the gene set is predominantly enriched in cancer, particularly Non-small cell lung cancer and Renal cell carcinoma. These two cancers share common genes in their prognostic gene lists, namely SYK, PLK1, and ADRA2C **(Figure S1)**. This implied that the CGS_ARCR_ may play a crucial role in the pathogenesis of kidney and lung cancers.

### 2.12. The comprehensive scoring system for potential indications

**Figure 12(A)** displays the ROC curves for the training and testing sets. The training set achieved an AUC value of 0.985, and the testing set achieved an AUC value of 0.967, demonstrating excellent performance in distinguishing between the indication and non-indication samples. The high AUC values suggest that the model possesses high sensitivity and specificity. The similar shapes of the ROC curves for the testing set compared to the training set demonstrate that the model maintains good generalization capability on the testing sets. **Figure 12(B)** illustrates the trend of log-loss for both the training and testing sets as the number of iterations increases. The log-loss decreases and stabilizes as iterations progress, indicating that the model gradually optimizes during iteration, and significantly reduces the loss. The log-loss decreases markedly for the training set during the initial iterations, followed closely by a similar trend in the testing set, leading to eventual convergence. This highlights the model’s robust stability and reliable convergence properties. **Figure 12(C)** shows the comparison between predicted values and actual training labels. The distribution of predicted values is depicted for samples with labels 0 and 1. Samples labeled as 0 exhibit predicted values concentrated in the low-value range, while samples labeled as 1 show predicted values predominantly in the high-value range. This distribution indicates the model effectively distinguishes between different classes and maintains high accuracy in predictions. **Figure 12(D)** presents the ROC curve at the optimal threshold, marked at 0.5689. This threshold is chosen to maximize the difference between sensitivity and specificity, ensuring the best classification performance, which demonstrates the model’s effectiveness and reliability in practical applications.

**Figure 12.**
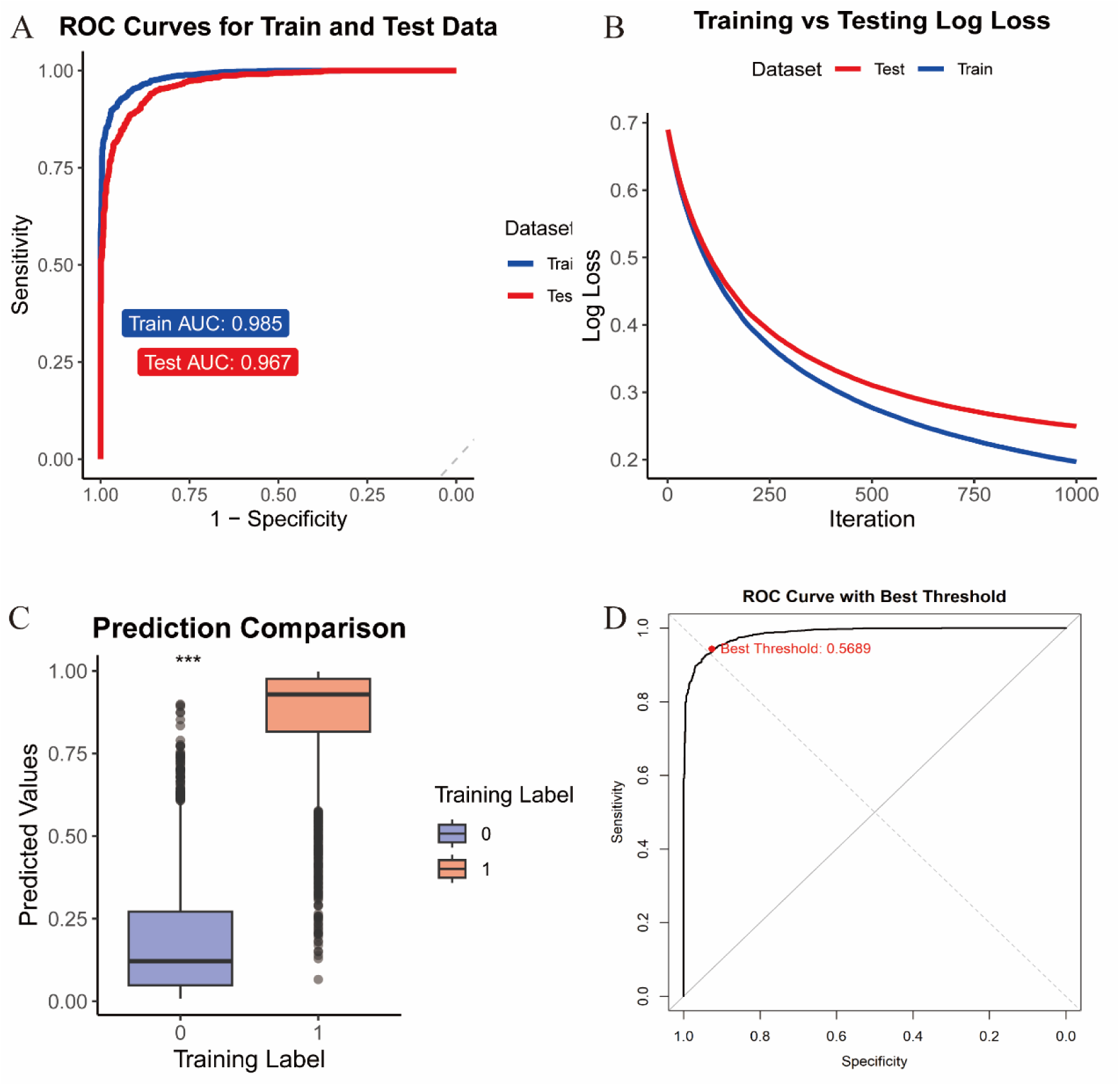
Performance Evaluation of Indications Prediction system based on XGboost. **(A)** ROC curves for the training and test datasets. **(B)** Log loss over iterations for the training and test datasets. **(C)** Comparison of predicted values with training labels. **(D)** ROC curve with the optimal threshold.

## 3. Discussion

Current pharmaceutical practice in drug repurposing primarily focuses on single-compound Western medicines for known diseases. Existing drugs, having undergone rigorous clinical evaluations to determine safety and pharmacokinetics, offer a reliable pathway for repurposing without the need for extensive clinical trials^10^. As existing drugs have undergone comprehensive clinical evaluations to assess safety and pharmacokinetics, their repurposing for novel indications offers a reliable pathway without the necessity for clinical experimentation, facilitating an investigation of their clinical therapeutic potential ^32^. However, the inherent heterogeneity of cancer initiation and progression poses significant challenges to the success of single-drug repurposing efforts, often yielding sporadic and limited outcomes ^14–16^. For example, while aspirin and metformin have demonstrated clinical efficacy as anti-cancer agents, they have not yet secured approval for treating advanced cancer patients, reflecting the complexities of cancer pharmacotherapy ^33,34^. The strategic optimization of drug repurposing in precision oncology, therefore, necessitates a synergistic combination of targeted drugs. TCM formulations, with their diverse array of natural compounds, offer a multi-component and multi-target approach to gene modulation, presenting a robust resource for refining drug repurposing strategies. The Astragali Radix-Curcumae Rhizoma (ARCR), a TCM formulation, has attracted considerable attention for its promising anti-tumor properties across various cancer types including lung, colorectal, gastric, and breast cancer. ARCR has shown efficacy in enhancing blood circulation and immunity within tumor lesions, leading to disease stabilization and symptomatic relief, and in some cases, disease regression—thereby contributing to extended survival for cancer patients ^10^.

The Astragali Radix-Curcumae Rhizoma (ARCR), a TCM formulation, has garnered considerable attention for its promising anti-tumor properties across various cancer types, encompassing lung cancer ^22,27^, colorectal cancer ^23,24,28^, gastric cancer ^29,35^, and breast cancer ^26^. Notably, ARCR has demonstrated efficacy in enhancing blood circulation and immunity within tumor lesions, resulting in disease stabilization, amelioration of symptoms, and especially instances of disease regression, thereby contributing to extended survival periods for cancer patients ^22–24,26–29,35^. These findings accentuate the remarkable anti-tumor potential intrinsic to ARCR. However, due to the complexity of TCM formulations, systematic research on the repurposing of these formulations in the field of cancer has been challenging, which has resulted in the underutilization of the potential therapeutic benefits of TCM in cancer treatment. Therefore, there is a clear need to develop a methodology for the reuse of TCM formulations, which takes full account of the complexity of their compounds and utilizes clinical data to systematically unravel the intricate connections between TCM and a spectrum of cancers.

In this study, we have developed a novel approach, NetPharm-PanCan, which combines network pharmacology and pan-cancer analysis. This approach is designed for the systematic identification of the representative core gene set associated with TCM formulations that exhibit broad-spectrum anti-tumor effects to further achieve their repurposing (**Figure 1**). In this regard, we utilize extensive data encompassing transcriptomic profiles of cancer patients, herb information, and relevant signaling pathways for the construction of CGS_ARCR_ to explore potential indications of ARCR in the field of cancer. Utilizing the Degree algorithm and the MCODE algorithm based on enrichment analysis, we identified a set of 28 genes within the CGS_ARCR_ that interact with the 27 components of ARCR. The CGS_ARCR_ can be effectively utilized to accurately predict the prognosis across seven potential cancers (**Table S20**). Among them, LUAD, COAD, BRCA, and STAD are well-studied indications of ARCR, while THCA, KIRC, and ESCA represent novel indications that have yet to be reported. Enrichment analyses of the CGS_ARCR_ and the prognostic gene sets revealed similar and distinct biological mechanisms involved in the prognostic markers across the seven potential indications (**Figure 11D**). Notably, KIRC and LUAD emerged as the two cancers most significantly associated with the CGS_ARCR_ (**Figure 11E**), demonstrating reliable and substantial predictive performance in their respective prognostic models (**Table S20**). Spearman correlation analysis indicated that the overexpression of the CGS_ARCR_ is not only associated with multiple cancer pathways but also linked to the infiltration of various immune cells, thereby exerting direct or indirect influences on the onset and progression of cancer (**Figure 8**). Finally, we established an indication prediction scoring system based on the XGboost algorithm to comprehensively evaluate the performance of NetPharm-PanCan. The results show that our method demonstrated a strong generalization ability for drug repurposing of ARCR. These findings demonstrated the effectiveness of NetPharm-PanCan in repurposing TCM formulations. In summary, NetPharm-PanCan provides a systematic approach for repurposing TCM formulations and if broadly applied, it holds promise for application in the field of cancer and other disease areas.

### 3.1. ARCR in previously studied cancers

Up to now, research on the ARCR formulation has been reported in LUAD, COAD, STAD, and BRCA. Notably, while studies focusing on ARCR in BRCA are sparse, with only one experimental investigation reported, comprehensive network pharmacology analyses and experimental validations in LUAD, COAD, and STAD substantiate ARCR therapeutic efficacy. These investigations have elucidated key genes and signaling pathways integral to ARCR’s effects in cancer treatment, as detailed in **Table 2**. Within these four cancer contexts, several critical genes identified in the literature, including AKT1, VEGFA, EGFR, PTGS2, and PIK3CA, have been found to overlap with our constructed CGS_ARCR_. These genes are confirmed to be upregulated specifically in these cancers, as presented in **Table 3**. Additionally, the collective expression levels of the CGS_ARCR_ exhibit a notable upregulation across these four cancers (**Figure 7A**), reinforcing the credibility of these genes as essential therapeutic targets and affirming the reliability of our constructed CGS_ARCR_. The signaling pathways linked with CGSARCR closely match those identified in previous studies on these four cancers, including the PI3K-Akt signaling pathway, VEGF signaling pathway, Ras signaling pathway, MAPK signaling pathway, and HIF-1 signaling pathway, as shown in **Figure S2**. These pathways are critical in our analysis, underscoring their importance in the therapeutic mechanisms of the ARCR formulation. The consistency of these findings across different data sources and methodologies emphasizes the robust potential of ARCR as a multifaceted therapeutic agent in oncology.

**Table 2.** Literature reports on the ARCR in four established cancer types.

| Cancer type | Bioinformatics | Clinic |
| --- | --- | --- |
| LUAD | Yang *et al.* <sup>27</sup> demonstrated that the key targets of ARCR in the treatment of lung cancer were VEGF, EGFR, and HIF-1α, and the compound inhibited angiogenesis in lung cancer by acting on the EGFR/PI3K/Akt, HIF-1, and MAPK signaling pathways. | Yang *et al.* <sup>36</sup> found that ARCR suppressed the expression of HIF-1α, EGFR, and VEGF, and this inhibition played a vital role in restraining lung cancer angiogenesis, as well as curbing the growth and metastasis of tumors. |
| COAD | Liu *et al.* <sup>28</sup> have demonstrated that the key targets were AKT1, PTGS2, VEGFA, etc. and the most relevant signaling pathways for the treatment of colorectal cancer by ARCR were Ras, PI3K/Akt and MAPK signaling pathways. | Wang AL *et al.* <sup>23</sup> demonstrated that formononetin from Astragalus inhibited colon cancer cell functions via PI3K/Akt and STAT3 pathways, achieving anti-tumor effects, while Wang JL *et al.* <sup>24</sup> showed that curcumol from Curcuma blocked cell growth in G0/G1 phase by inactivating Akt/GSK3β/cyclin D1 signaling pathways. |
| STAD | Huang *et al.* <sup>29</sup> highlighted that TP53, MYC, CASP3, AKT1, and JUN were the main targets of ARCR for gastric cancer treatment and the key pathways were the PI3K/Akt and MAPK signaling pathways | Meng *et al.* <sup>25</sup> showed that ARCR inhibited the progression of gastric cancer by suppressing the expression of Akt and Akt signaling pathway, thereby reducing cell proliferation and promoting apoptosis in gastric cancer MGC803 cells. |
| BRCA | — | Deng *et al.* <sup>26</sup> demonstrated that ARCR achieved inhibition of breast cancer through the downregulation of PI3K, Akt1, and Akt2 as well as the upregulation of PTEN gene expression by establishing a nude mouse transplantation tumor model of human breast cancer cell line. |

**Note:** The sign — Indicates the absence of studies on the ARCR in that specific cancer field.’

**Table 3.** Reports on Key Genes Overlapping Between the CGS_ARCR_ and Previous Studies.

| Gene | Cancer type | Abstract |
| --- | --- | --- |
| AKT1 | LUAD | Carpten *et al.* <sup>37</sup> showed that AKT1 was highly expressed in lung cancer. Yang *et al.* <sup>27</sup> demonstrated that ARCR reduced PI3K levels by inhibiting its phosphorylation process, consequently leading to the downregulation of AKT and inhibiting AKT protein production. |
|  | STAD | Carpten *et al.* <sup>37</sup> demonstrated that AKT1 was highly expressed in gastric cancer, involving the occurrence, development, and infiltration of gastric cancer. Meng *et al.* <sup>25</sup> showed that ARCR inhibited Akt protein expression to obstruct the development of gastric cancer. |
| | COAD | Roy *et al.* <sup>38</sup> demonstrated that in colorectal cancer, AKT overexpression was recently described to be an early event during sporadic carcinogenesis. Wang *et al.* <sup>28</sup> found that ARCR leads to inactivation of the Akt/GSK3β/cyclin D1 signaling pathway. |
|  | BRCA | Hutchinson *et al.* <sup>39</sup> pointed out that the overexpression of Akt1 in the mammary epithelial cells inhibited the pro-apoptotic signals, while activating the survival signals to support the process of tumorigenesis. Deng *et al.* <sup>26</sup> demonstrated that ARCR achieved inhibition of breast cancer through the downregulation of Akt1. |
| VEGFA | LUAD | Kajdaniuk *et al.* <sup>40</sup> found that VEGF overexpression and/or high VEGF serum levels have been confirmed in non-small-cell lung carcinoma (NSCLC) and small-cell lung carcinoma (SCLC). Yang *et al.* <sup>27</sup> demonstrated that ARCR significantly inhibited the expression of VEGF, thereby inhibiting tumor angiogenesis |
|  | COAD | Lee *et al.* <sup>41</sup> found that VEGF was highly expressed in Colorectal cancer. |
| | STAD | Natsume *et al.* <sup>42</sup> reported that VEGFA in gastric cancer cells was also upregulated through HIF-1α upregulation, resulting in angiogenesis. |
| EGFR | LUAD | Normanno *et al.* <sup>43</sup> found that the prognostic significance of EGFR overexpression occurred in approximately 40–80% of NSCLC patients. Yang *et al.* <sup>27</sup> found that ARCR suppressed VEGF expression, thereby inhibiting angiogenesis, tumor growth, and metastasis in lung cancer. |
|  | STAD | Dulak *et al.* <sup>44</sup> reported that EGFR was overexpressed in 27%–64% of gastric tumors and its role as an oncogene in this malignancy was well-known. |
| PTGS2 | LUAD | A direct relationship between COX-2 overexpression and the poor survival of patients with lung carcinoma has been demonstrated in the study of Khuri <sup>45</sup>. |
|  | COAD | Eberhart *et al.* <sup>46</sup> indicated that the overexpression of COX-2 was associated with various premalignant and malignant lesions of epithelial origin, in gastrointestinal tract organs. |
| PIK3CA | LUAD | Angulo *et al.* <sup>47</sup> pointed out that PIK3CA gene amplification in lung SCCs was common and was correlated with overexpression. |

Upon further exploration, we learned AKT1 was a common key target of the ARCR in the studies of these four cancers (**Table 3**). AKT1 serves as a critical regulatory factor for cell survival and proliferation, and its upregulation or activation has been associated with tumor growth, metastasis, and increased resistance to cancer treatment ^48^. The PI3K-Akt signaling pathway, featuring the involvement of AKT1, stands out as a pivotal common pathway in these four cancers (**Table 3**). This pathway was identified as one of the central regulators modulated by the ARCR for the treatment of diverse cancers (**Figure S2**), as extensively validated in clinical experiments (**Table 2**). Concurrently, the enrichment analysis of the CGS_ARCR_ underscored a pronounced enrichment of the PI3K-Akt-signaling pathway (**Figure 11E**), reinforcing the correlation between this pathway and ARCR. Lawrence *et al.* ^49^ indicated that the PI3K-Akt-signaling pathway is implicated in the onset of various cancers, comprising breast cancer and lung cancer. Corresponding drugs, such as PI3K and AKT inhibitors, have been developed to interfere with this pathway for cancer treatment ^50^. Notably, compounds within ARCR, including Quercetin and Kaempferol from Astragali Radix (AR) and Wenjine from Curcumae Rhizoma (CR), show potential as AKT1 inhibitors (Table S7). Studies indicate that low doses of Quercetin can inhibit several enzymes involved in cell proliferation and signaling pathways in lung and breast cancer cells ^51^.

The consistency observed across these findings reinforces that CGSARCR, constructed using NetPharm-PanCan, accurately represents ARCR functional significance in oncology. This validates the computational results’ reliability and demonstrates NetPharm-PanCan potential to explore the broader application of TCM formulations in cancer treatment.

Beyond corroborating previous findings, our study reveals novel insights into the four established indications. By integrating pharmacology and genetic data, we constructed a comprehensive CGS_ARCR_ that intricately interacts with all 27 ARCR components (**Figure 6C**). This gene set provides a more detailed portrayal of ARCR pharmacological characteristics compared to previous studies. Furthermore, prognostic models built on CGSARCR exhibited robust predictive performance across the four cancers, with varying reliability over different time intervals, suggesting that the therapeutic efficacy of ARCR may differ across cancer types. This variation likely reflects ARCR distinct impact on the pathological features of each cancer, contributing to different prognoses among patients.

In particular, our model for LUAD demonstrated higher predictive reliability compared to models for other cancers. Enrichment analysis further highlighted a strong correlation between CGS_ARCR_ and LUAD, especially with the HIF-1 signaling pathway (**Figure 11E**). This finding aligns with Xu et al.^22^ on LUAD indicated that the ARCR reduce tumor micro vessel density, inhibit VEGF expression, and suppress the TGF-β1 and HIF-1α expression, exerting inhibitory effects on Lewis’s lung cancer tumor growth and metastasis. These results underscore the significant therapeutic potential of ARCR for LUAD. The CGS_ARCR_ also demonstrates a robust correlation with the distinctive biological processes of LUAD, implying a multifaceted inhibition of cancer development through various pathways. Moreover, the enrichment analysis of the prognosis gene set of LUAD revealed that the most significant signaling pathway is the G2/M transition of the mitotic cell cycle (**Figure 11D**). Aberrations in this pathway can lead to uncontrolled cell division, promoting tumor growth and spread ^52^. This implied that the components of ARCR associated with the prognosis genes of the LUAD model may hinder the progression of lung cancer through the regulation of certain cell cycle-related prognosis genes, such as mutations or dysregulation of WEE1 ^53^ and CDC25B ^54^ (**Table S21**). This hindrance may impede lung cancer cells from entering the G2/M transition phase of the cell cycle, inhibiting uncontrolled proliferation and division, which could potentially contribute to therapeutic effects (**Table S21**). It is noteworthy that current research on the therapeutic effects of the ARCR in lung cancer has predominantly focused on the LUAD subtype, with limited exploration into LUSC, another subtype of lung cancer. ARCR may have limited therapeutic effects on LUSC, due to the suboptimal predictive ability of its prognostic model constructed in this study (**Figure 9I**). This may explain the lack of ARCR research on LUSC in the literature.

In the models of COAD, BRCA, and STAD, some identified prognosis genes have been validated to be overexpressed in specific cancer tissues. Additionally, ABL1 and ESR1 in COAD, as well as ADRA2C, AR in BRCA, and ADRA2C, ESR1and NR3C1 in STAD, stand out as newly discovered prognostic markers in this study. The expression profiles of these genes in cancer remain unreported, necessitating further research to confirm their specific biological significance in the respective cancers (**Table S22-S24**).

Furthermore, our investigation reveals a close association between the CGS_ARCR_ and the immune microenvironment across the four cancer types, suggesting that the therapeutic effects of ARCR in cancer treatment extend beyond the direct inhibition of tumor cell growth. It may also involve modulating the activity of the immune system to achieve therapeutic outcomes, aligning with the current trend of immunotherapy in cancer treatment. The collective expression levels of the CGS_ARCR_ across the four cancers show a positive correlation with Th1 cells, DC cells, and iTreg cells (**Figure 8 B**). Th1 cells are typically associated with pro-inflammatory immune responses ^55^. While iTreg cells are essential for preserving immunological self-tolerance, their role involves actively suppressing self-reactive lymphocytes ^56^. DC cells play a pivotal role in initiating immune responses ^57^. Additionally, the balanced presence of immune cells contributes to maintaining the stability of the immune system while effectively combating tumor cells ^58^. So, it is essential to consider the role of immune cells showing a negative correlation with the CGS_ARCR_ expression such as CD8 T, MAIT, Gamma-delta, NK, and CD4 T cells, which might contribute to the overall immune regulation in the tumor microenvironment. Therefore, we hypothesized that ARCR may regulate the activity of different immune cells by targeting CGS_ARCR_ to maintain the stability of the immune system in cancer tissues and achieving anti-tumor effects. These findings support the concept that the ARCR acts against cancer by promoting immune responses, presenting promising prospects for future cancer treatment.

### 3.2. ARCR in Newly Discovered Cancers

In this study, our approach not only reproduced the four reported indications but also unveiled three novel indications: THCA, KIRC, and ESCA.

The pan-cancer analysis revealed a notable increase in the overall expression levels of the CGS_ARCR_ in these three types of cancer (**Figure 7 A**). The prognostic models for these cancers demonstrated reliable predictive capabilities, with THCA exhibiting the most reliable prognostic model among all cancers, KIRC’s model performing comparably to LUAD, and ESCA’s model showing performance similar to COAD, indicating that their prognostic abilities rank as THCA > KIRC > ESCA (**Table S20**). However, because of THCA’s high cure rate of 80-90%^59^, the positive prognosis may lead to its high performance of prognostic model. In contrast, applying the ARCR to KIRC holds substantial promise. This is because that KIRC is experiencing an annual increase in occurrence, ranking among the top 10 fatal cancers in patients^60,61^ and the prognostic model based on ARCR demonstrates high reliability for KIRC, supported by the enrichment analysis of CGS_ARCR_ indicating a significant association with renal cell carcinoma (**Figure 11E**). These findings underscore the potential of ARCR in treating KIRC effectively compared to THCA. Furthermore, studies on the prognosis-related genes for the three new indications also indicated that certain prognosis genes can act on the progression of the corresponding cancers (**Table S25-27**). Within the prognostic gene set for KIRC, PLK1 ^62^, SRC ^63^, and AR ^64^ have been shown to be highly expressed in the tissue of renal clear cell carcinoma, leading to poor prognosis. These genes facilitate the invasion, migration, and proliferation of renal clear cell carcinoma cells (**Table S25**). In the ESCA prognosis model, four prognosis genes, including CDC25A ^65^, CDC25B ^66^, CHEK1 ^67^, AKT1 ^68^, have been demonstrated to be overexpressed in tumor tissues, resulting in an unfavorable prognosis (**Table S26**). Similarly, Among the 13 prognosis genes in THCA, six genes, including CDC25A ^69^, CDC25B ^70^, APP ^70^, RAF1 ^71^, ESR1 ^63^, and PIK3CA ^72^, have been individually confirmed to exhibit high expression levels in tumor tissues, leading to poor prognosis (**Table S27**). These genes may be crucial for the therapeutic effects of the ARCR on new indications, and the intricate cancer-related mechanisms related to them hold promise as prospective targets for the ARCR in the realm of cancer treatment.

During the process of drug repurposing, a noteworthy discovery is the positive correlation between the CGS_ARCR_ and Cell cycle pathway in 20 out of 33 cancer types, encompassing the seven indications under our investigation (**Figure 8A**). Furthermore, the enrichment analysis of the CGS_ARCR_ revealed the highest significance in the cell cycle pathway within cellular processes (**Figure 11E**), which has not been thoroughly explored in previous studies. The cell cycle, as evidenced by prior research ^73^, emerges as a pivotal pathway frequently disrupted in diverse cancers. The regulatory factors of cell cycle serve as downstream targets of crucial pathways such as PI3K-AKT, and their alterations intertwine with the progression of cancer ^74^. This indicates a likelihood that components within the ARCR exert their influence on the regulatory factors of cell cycle, including PLK1, ABL1, CDK2, housed within the CGS_ARCR_ to inhibit the progression of cell cycle pathway in cancer, impeding the cancer cells proliferation. These findings represent crucial mechanisms through which ARCR could effectively treat lung, gastric, colorectal, and breast cancers, potentially paving the way for drug repurposing strategies. In addition, cell cycle regulators were also prevalent in the prognosis genes of the three new indications, comprising notable candidates such as PLK1, CDC25A, CDC25B, and CHEK1 (**Table S24-26**). This observation underscores the potential pivotal role of regulatory factors of the cell cycle in cancer prognosis, thereby suggesting a broad prospect for their development as biomarkers across various cancers. Not to be overlooked, the yet-unreported prognostic genes, such as the shared HSP90AA1 common to the three cancers and the shared PTGS2, APP, ADRA2C in KIRC and THCA (**Table S24-26**), have the potential to serve as biomarkers for the potential indications. These insights open avenues for the exploration and development of novel biomarkers with broad applications in cancer prognostication.

### 3.3. Similarities and Differences among the Seven Indications of the ARCR

Enrichment analysis of the prognostic genes across seven indications revealed common involvement in biological processes, specifically kinase binding and positive regulation of phosphorylation (**Figure 11D**). These processes were intricately linked to the PI3K/Akt-signaling pathway, where AKT and PI3K act as kinases, interacting with other proteins in the signal transduction process ^75^. Moreover, Elevated kinase activity is crucial for fostering Akt phosphorylation, activation, and ultimately, pathway stimulation ^76,77^. These results proposed that the PI3K/Akt-signaling pathway emerges as vital for ARCR efficacy, not only in treating known indications but also for new ones. Similar to the cell cycle, this pathway may lead to the broad-spectrum anti-cancer therapeutic effects of TCM.

Furthermore, we identified some key regulatory factors, including AKT1, PLK1, EGFR, CDK, VEGFA, etc. (**Figure 6C**). These factors serve as both prognostic genes and crucial regulatory genes in the PI3K/Akt-signaling pathway or cell cycle pathway. By downregulating the expression of these genes, the ARCR may exert a synergistic effect on multiple targets and pathways to inhibit cell proliferation, migration, and invasion of cancer, addressing the heterogeneity of different cancers from various perspectives to achieve effective therapeutic outcomes.

The mechanisms of action for the ARCR on established and novel indications exhibit both commonalities and differences. The Commonalities are mainly observed in the mechanisms targeting cancer (**Figure 11D**). The differences, on the other hand, are primarily reflected in the varying effect of key prognostic genes for different cancers (**Table S19**), which leads to further distinctions in the biological characteristics of different cancer types. For instance, PLK1 acts as a risk gene (HR > 1) in BRCA (**S11**) and LUAD (**Table S12**) but as a protective gene (HR < 1) in COAD. (**Table S14**)

Overall, the mechanisms of action for ARCR on established and novel indications exhibit both commonalities and differences. The commonalities are mainly observed in the mechanisms targeting cancer through the PI3K/Akt-signaling pathway and cell cycle regulation (**Figure 11D**). The differences are primarily reflected in the varying effects of key prognostic genes across different cancers ().

Future research on the ARCR should focus on experimentally confirming the clinical therapeutic effects on the three novel indications and applying them to cancer treatment. Additionally, attention should also be directed towards understanding the roles of the PI3K/Akt-signaling pathway and cell cycle in both established and novel indications. This investigation will contribute to a deeper understanding of the molecular mechanisms underlying the potential indications of ARCR, clarify the genetic regulatory basis, and provide insights into immune therapy strategies.

### 3.4. Innovation and Limitations

The innovation of this study lies in both its methodology and application. Methodologically, we present a novel approach by combining network pharmacology, pan-cancer analysis, and prognostic modeling to reveal the associations between drugs, targets, and diseases. This integrated methodology allows for the precise repurposing of Traditional Chinese Medicine (TCM). Unlike existing methods that primarily rely on bioinformatics analysis of single gene targets, our study expands the focus to examine the collective effects of gene sets. We assess the role of differentially expressed target sets across various cancers, rather than targeting a single key gene. This holistic approach enables the prediction of potential new indications for drugs at a more comprehensive level.

Our study emphasizes the reliability of drug repurposing by validating the key genes and mechanisms involved in the studied indications. Additionally, based on the expression levels of core gene sets across all samples and indications, we employed the XGBoost algorithm to construct a system capable of identifying potential new indications for drugs.

In terms of application, this study marks the first attempt to reposition TCM, using ARCR as an example. Our results, which align with previous research, reaffirm the efficacy of four existing indications and identify three new ones, demonstrating the potential of our method in TCM repurposing. Furthermore, our research explores the mechanisms behind “same disease, different treatment” and “different disease, same treatment,” shedding light on the multifunctional and multi-pathway therapeutic effects of TCM compounds in diverse disease contexts. Thus, this innovative approach may significantly expand the applicability of TCM formulations and enhance their potential in cancer treatment.

However, our study does have some limitations. First, our methodology does not account for the dosage of ARCR components in the context of drug repurposing. Additionally, as our research relies on existing data, the quality and credibility of the information used is crucial, and further validation is needed.

## 4. Method

### 4.1. Data collection

In this research, the primary active constituents of ARCR were obtained through the utilization of the online platform TCMSP (https://tcmsp-e.com/load_intro.php?id=43) with the keywords “Astragalus (Huangqi)” and “Curcuma longa (Ezhu)”. All components were subjected to activity screening by absorption, distribution, metabolism, and excretion (ADME) related parameters according to the criteria of drug-likeness (DL) greater than or equal to 0.18 and oral bioavailability (OB) greater than or equal to 30%. Additionally, a meticulous examination of pertinent literature was conducted for the supplement of the principal bioactive components inherent in both TCMs.

To perform pan-cancer analysis, overall 11,093 samples from 33 different types of cancer were obtained, including mRNA expression profile data (n = 10,471) and clinical data (n = 11,160) for these cancer types, from the TCGA database (https://www.tcpaportal.org/tcpa/) and the Gene Set Cancer Analysis database (GSCA, http://bioinfo.life.hust.edu.cn/GSCA). Moreover, RPPA data were obtained from the TCPA database (https://www.tcpaportal.org/tcpa/).

### 4.2. Target prediction of ARCR

The ARCR components’ data in SMILES format was obtained utilizing the PubChem database (https://pubchem.ncbi.nlm.nih.gov/). The Swiss Target Prediction database (http://www.SwissTargetPrediction.ch/) was employed to predict targets associated with each active components of the ARCR. The target genes were corrected using the Uniprot database (https://www.uniprot.org/), and the species was limited to “Homo sapiens” to convert the target proteins into the corresponding genes.

### 4.3. Topology analysis and construction of PPI networks

The ARCR-component-target gene network was constructed and subjected to topological analysis using Cytoscape (version 3.7.2). Topological analysis was performed using Cytoscape’s network analysis tools, with the median degree value of target genes serving as the cutoff for preliminary gene screening. The identified target genes were then uploaded to the STRING database (https://string-db.org/) for further interaction analysis. A Minimum Required Interaction Score of 0.4 and species restriction to “Homo sapiens” were applied, and genes with no interactions were removed, resulting in a highly interconnected PPI network.

### 4.4 Construction of the core gene set of ARCR

The genes from the PPI network were input into the Metascape database (https://metascape.org/) for Gene Ontology (GO) and Kyoto Encyclopedia of Genes and Genomes (KEGG) functional enrichment analysis. The results were visualized using circos plots. Subsequently, a protein function enrichment network was created using the BioGrid database (https://thebiogrid.org/), InWeb_IM and OmniPath8 plugins. This network extracted protein interactions among all target genes from the submitted PPI network and assigned biological significance to these genes based on the results of KEGG enrichment analysis. The MCODE algorithm in the Metascape database was applied to cluster this functional network, and the subclass of highest scores with the most genes was extracted, which was defined as the core functional gene set of ARCR. Simultaneously, the Degree algorithm from the Cytohubba tool in Cytoscape was applied to analyze the target genes in the PPI network and the top ten genes with the highest degree values were selected as the core hub gene set of ARCR. Finally, these two sets were merged to form the core gene set of ARCR (CGS_ARCR_) after the duplicates were removed.

### 4.5. Differential analysis of GSVA in normal and tumor samples

The Gene Set Variation Analysis (GSVA), conducted in an unsupervised manner, was able to calculate the GSVA scores of the core gene set in both tumor and normal samples. These scores reflect the comprehensive expression level of the inputted gene set, displaying a positive correlation with its expression. In this study, GSVA was conducted to assess the differences in expression levels of the CGS_ARCR_ between tumor and normal samples for exploring the potential indications of ARCR on the condition that the cancer types had more than 10 pairs of normal-tumor sample. The GSVA scores were computed using the R package GSVA, and the results were visualized through box plots. When the GSVA scores of specific tumor samples exceeded those of normal samples, i.e., log2 FC (T/N) > 0 and *P* < 0.05, it was deemed that the CGS_ARCR_ exhibited statistically significant overexpression in that tumor.

### 4.6. GSEA analysis

The Gene Set Enrichment Analysis (GSEA) was carried out on the latent indications obtained from GSVA differential analysis using the R package fgsea (46). Differential gene expression analysis was conducted on all genes in the pan-cancer genome (N = 20,000), assessing the differential expression of each gene and ranking them based on gene expression fold change. Subsequently, the Enrichment analysis was employed to evaluate whether genes of the CGS_ARCR_ were significantly enriched in specific cancers compared to all genes. The Enrichment Score (ES) reflected the collective enrichment of the core gene set relative to the entire genome. A positive ES signified a noteworthy enrichment of genes within the set in a specific cancer type, whereas a negative ES indicated a lack of enrichment under the same context. The results were visualized through enrichment plots, illustrating the relative enrichment of CGS_ARCR_ compared to all genes.

### 4.7. Correlation of GSVA scores with cancer-related pathways

The correlation between GSVA scores of CGS_ARCR_ and cancer-related pathways was assessed based on pathway activity scores which were computed by subtracting the sum of protein levels of all negative regulatory components from the protein levels of all positive regulatory components within pathways. The RPPA data from the TCPA database were used to calculate relative protein levels for pathway activity analysis. Specifically, pathway activity scoring was applied to ten renowned cancer-related pathways, including TSC/mTOR, RTK, RAS/MAPK, PI3K/AKT, hormone ER, hormone AR, EMT, DNA damage response, cell cycle, and apoptosis pathways. These scores were assigned to evaluate the relationship between the comprehensive expression levels of the CGS_ARCR_ and the activity of these cancer-related pathways across 33 cancer types by Spearman correlation analysis, with *P* ≤ 0.05 and FDR ≤ 0.05 signifying significant correlations. The results were visualized using a heatmap generated by “ggplot2” package.

### 4.8. Correlation of GSVA score with immune infiltration

The ImmuCellAI tool (http://bioinfo.life.hust.edu.cn/ImmuCellAI<u>#!/</u>) was used to evaluate the infiltration levels of 24 different kinds of immune cells. The association between immune infiltration and CGS_ARCR_ GSVA scores was investigated using the Spearman correlation analysis, where the coefficient denotes the strength of the interaction between immune cells and CGS_ARCR_ collective expression levels with *P* ≤ 0.05 and FDR ≤ 0.05 signifying significant correlations. Visualization of the results was achieved using a heatmap generated by “ggplot2” package.

### 4.9. Construction of prognostic model

The mRNA expression profiling data and clinical data of cancers downloaded by TCGA were used to perform multivariate Cox regression analysis on the genes of the CGS_ARCR_ to identify the prognostically relevant signature genes and calculate the *P*-value, coefficient, HR, and 95% CI for each of the prognostic genes. The prognostic models were constructed in accordance with the following equation:

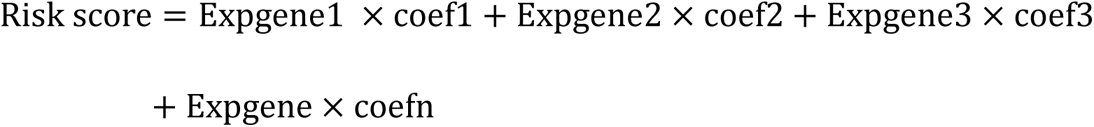

Where Expgene represents the gene expression levels and coef represents the risk coefficient of the corresponding gene. Each gene has a risk coefficient (coef), where negative values indicate protective genes and positive values indicate risk genes. Based on the median risk score, patients were categorized into high-risk and low-risk groups. The model’s performance was evaluated using Kaplan-Meier survival analysis and receiver operating characteristic (ROC) curves. Kaplan-Meier curve plotting, log-rank tests, and Cox proportional hazards regression analysis were conducted using the R packages “survival” and “survminer.” ROC curves were generated using the R package “pROC,” with the area under the curve (AUC) and confidence intervals assessed using the ci function of pROC to obtain the final AUC result.

### 4.10. Analysis of prognostic genes and CGS_ARCR_

Metascape (https://metascape.org/gp/index.html#/main/step1) was utilized to analyze and identify shared or selectively attributed pathways and biological processes among specific gene lists from reliable prognostic models. This involved constructing networks to unveil common prognostic genes and employing enrichment analysis to discern variations in prognostic genes across different cancer types. Leveraging the most recent gene annotations for KEGG Pathways obtained via the KEGG rest API (https://www.kegg.jp/kegg/rest/keggapi.html) as a background, all genes of the CGS_ARCR_ were mapped for further enrichment analysis using the clusterProfiler R package. Circos plots were used to illustrate similarities of gene composition and pathway among the prognostic gene lists, and pie charts were employed to clarify common genes among these lists. Cluster heatmaps were employed to depict the most enriched clusters and variations in enrichment patterns of GO biological process and KEGG signal pathways across multiple gene lists. The KEGG enrichment pathways of the CGS_ARCR_ were further presented through bar graphs.

### 4.11. Comprehensive scoring system of indications

Given that our drug repurposing approach is based on the overall expression of gene sets, the system we constructed using machine learning should be capable of determining whether a particular disease is an indication for the drug based on the overall expression of core genes. The mRNA expression levels of 28 genes from the CGS_ARCR_ across 14 types of cancer were extracted from the TCGA database (**Supplementary Sheet 5**). Based on prior analyses, all samples for each cancer type were labeled as either True or False, with True indicating a potential indication for the drug, and False indicating a non-indication. The expression data with labels were used to construct the dataset for model training. The dataset was randomly divided into training and testing sets with 60% and 40% of all samples, respectively. Features and labels were extracted from both sets and converted into numerical values to fit the input requirements of the XGBoost model.

The XGBoost algorithm was applied to create the model, with several key parameters set to ensure robustness and high performance. These parameters included a binary logistic regression objective function, log-loss evaluation metrics, a low learning rate, shallow tree depth, and high sampling ratios for both data and features. Enhanced L1 and L2 regularization parameters were also implemented to further prevent overfitting. Optimal parameters and the number of training iterations were selected through cross-validation, and early stopping techniques were employed to optimize the model. The feature matrix was then imported into the trained model for classification predictions.

The model’s ability to distinguish between indications and non-indications was evaluated using the area under the ROC curve (AUC). To find the optimal balance point, the coords () function was utilized, which automatically identifies the best compromise between sensitivity and 1-specificity (false positive rate). This point represents the optimal threshold for determining whether a condition is an indication of the ARCR drug system.

### 4.12. Statistical analysis

All data processing was carried out via R software (v4.3.1) for statistical analysis. Spearman correlation analysis was conducted using the “cor.test()” function within the “stats” package in R for correlation analysis. Multivariate Cox regression analysis was performed using the “coxph()” function within the “survival” package in R to calculate survival risk and HR. ROC curves of prognostic models were generated using the “pROC” package in R, and their AUC values were calculated using the “roc()” function. Significance for Kaplan-Meier curves was assessed using log-rank tests and *P* < 0.05 was considered statistically significant. In this study, all data were presented as mean ± standard error of the mean (SEM), and group differences were analyzed using t-tests.

## 5. Conclusion

In summary, we developed a systematic drug repurposing approach, NetPharm-PanCan, to investigate the antitumor potential and explore potential indications for traditional Chinese medicine (TCM) in the context of ARCR. This method provides an empirical framework for repurposing TCM and highlights the significant repurposing potential of herbal medicines. By integrating network pharmacology and pan-cancer analysis, we were able to validate key genes and mechanisms for four established indications while also identifying three previously unreported indications of ARCR, including KIRC, ESCA, and THCA, across various aspects. Additionally, we discovered novel prognostic genes and elucidated their associated immune and pathway mechanisms for the established indications. The prognostic models developed for these seven potential indications offer strong support for both existing and new use cases, thereby reinforcing the robustness of our findings and methodology. Our research establishes a solid foundation for future investigations into the repurposing of TCM from a novel perspective.

## Supporting information

Supplemental Sheet 1-5

## Abbreviations

TCM: Traditional Chinese Medicine
NetPharm-PanCan: Network pharmacology and pan-cancer analysis
ARCR: Astragali Radix-Curcumae Rhizoma
WGS: whole genome sequencing
TCMSP: Traditional Chinese Medicine Systems Pharmacology
TCGA: The Cancer Genome Atlas
GSCA: Gene Set Cancer Analysis database
RPPA: Reverse phase protein array
PPI: Protein-Protein Interaction
GO: Gene Ontology
KEGG: Kyoto Encyclopedia of Genes and Genomes
CGS_ARCR_: Core gene set of ARCR
GSVA: Gene Set Variation Analysis
GSEA: Gene Set Enrichment Analysis
TCPA: The Cancer Proteome Atlas
ROC: Receiver perating characteristic
AUC: Area under curve
SEM: Standard error of the mean

## Declarations

### Ethics declarations

#### Ethics approval and consent to participate

Not applicable.

#### Consent for publication

Not applicable.

### Authors’ contributions

ZH, DY, YL (iu) and YL (iao) conceived and designed the study. DY collected the data, performed the analysis, interpreted the results, prepared all the pictures and tables, wrote, and revised the manuscript. DY, HX, AF and MS wrote the manuscript. LW revised the pictures and tables. ZH, WK and DO guided the writing of the manuscript, revised the manuscript and provided financial support. Additionally, all the authors read and approved the final manuscript.

### Availability of data and materials

The paper contains all necessary data to assess the presented conclusions, and supplementary materials also contribute to this information. Computation utilized existing code, and references or links to the code are provided in each respective method subsection.

## Acknowledgments

We thank Zhuang Kai for his technical expertise and support during the early stages of the research.

## Supplementary Information

**Additional file 1. Supplementary Table S1-S27, Figure S1-S7 and Supplementary sheet 1-6. Table S1.** The abbreviation of 33 types of cancer. **Table S2** Topological analysis of 442 targets in the ARCR -component-target gene network. **Table S3.** Gene of PPI network. **Table S4** Total score and composition of pathway enrichment in 8 subcategories. **Table S5** The top 3 KEGG Pathways of 8 Subclasses Classified by MCODE Algorithm. **Table S6** Genes of CGSARCR and their protein names. **Table S7** Genes of CGSOARCR and Corresponding Components of ARCR. **Table S8** Differential GSVA. **Table S9-S18** GSEA. Table S10. Results of univariate Cox regression analysis of CGSOA-CC in THCA, BRCA, KRIC, COAD, STAD, HNSC, ESCA, LUSC. **Table S19** Presence of 28 genes from the CGSARCR in seven potential indications. **Table S20** Predictive Performance Ratings of Prognostic Models for Nine Potential Indications. **Table S21-S27** Literature Reports on Prognostic Genes of ARCR in LUAD, COAD, BRCA, STAD, KIRC, ESCA, THCA. **Fig. S1** ARCR-component-target gene network. **Fig. S2** PPI network of ARCR. **Fig. S3** GO items and KEGG pathways sorted by P-value in the PPI network of ARCR. **Fig. S4** Survival curves of CGSARCR in potential prognostic models for nine different cancers. **Fig. S5** Overall Scores for Seven Potential Indications. **Figure S6** Intersection Genes in the Prognostic Gene Lists of LUAD and KIRC. **Figure S7** Network of pathway similarities and differences between CGSARCR and studies of LUAD, COAD, STAD, BRCA. **Supplementary sheet 1-4** The terms of GO-BP, GO-CC, GO-MF and KEGG. **Supplementary sheet 5** Data for normalization. **Supplementary sheet 6** Normalized data and their weighted average overall scores.

