## Supplemental Table S1-S27 and Figure S1-S2 for "Synergizing Network Pharmacology and Pan-Cancer Analysis in TCM Repositioning for Tumor Therapy"

**Table S1**. The abbreviation of 33 types of cancer

| **Cancer Type** | **Abbreviation** | **Number of Samples** | **Tumor-Normal Samples** |
| --- | --- | --- | --- |
| Adrenocortical carcinoma | ACC | 92 | 0 |
| Bladder Urothelial Carcinoma | BLCA | 411 | 19 |
| Breast invasive carcinoma | BRCA | 1,218 | 112 |
| Cervical squamous cell carcinoma and  endocervical adenocarcinoma | CESC | 310 | 3 |
| Cholangiocarcinoma | CHOL | 45 | 9 |
| Colon adenocarcinoma | COAD | 329 | 41 |
| Lymphoid Neoplasm Diffuse Large  B-cell Lymphoma | DLBC | 48 | 0 |
| Esophageal carcinoma | ESCA | 198 | 8 |
| Glioblastoma multiforme | GBM | 174 | 0 |
| Head and Neck squamous cell carcinoma | HNSC | 566 | 43 |
| Kidney Chromophobe | KICH | 91 | 23 |
| Kidney renal clear cell carcinoma | KIRC | 606 | 72 |
| Kidney renal papillary cell carcinoma | KIRP | 323 | 31 |
| Acute Myeloid Leukemia | LAML | 173 | 0 |
| Brain Lower Grade Glioma | LGG | 534 | 0 |
| Liver hepatocellular carcinoma | LIHC | 359 | 50 |
| Lung adenocarcinoma | LUAD | 576 | 57 |
| Lung squamous cell carcinoma | LUSC | 554 | 49 |
| Mesothelioma | MESO | 87 | 0 |
| Ovarian serous cystadenocarcinoma | OV | 309 | 0 |
| Pancreatic adenocarcinoma | PAAD | 198 | 4 |
| Pheochromocytoma and Paraganglioma | PCPG | 187 | 3 |
| Prostate adenocarcinoma | PRAD | 550 | 52 |
| Rectum adenocarcinoma | READ | 105 | 9 |
| Sarcoma | SARC | 265 | 2 |
| Skin Cutaneous Melanoma | SKCM | 474 | 1 |
| Stomach adenocarcinoma | STAD | 450 | 27 |
| Testicular Germ Cell Tumors | TGCT | 156 | 0 |
| Thyroid carcinoma | THCA | 122 | 58 |
| Thymoma | THYM | 572 | 2 |
| Uterine Corpus Endometrial Carcinoma | UCEC | 57 | 23 |
| Uterine Carcinosarcoma | UCS | 201 | 0 |
| Uveal Melanoma | UVM | 80 | 0 |

**Table S2** Topological analysis of 442 targets in the ARCR -component-target gene network

| Gene name | Degree | Betweenness Centrality | Closeness Centrality |
| --- | --- | --- | --- |
| EGFR | 10 | 0.021136 | 0.386831 |
| PTGS1 | 10 | 0.025687 | 0.362375 |
| CYP19A1 | 10 | 0.027163 | 0.391341 |
| HSD17B2 | 9 | 0.021956 | 0.34789 |
| CDK1 | 8 | 0.030441 | 0.376301 |
| ALOX12 | 8 | 0.013709 | 0.360153 |
| ALOX5 | 8 | 0.023218 | 0.378117 |
| ABCB1 | 8 | 0.013384 | 0.316498 |
| TYR | 7 | 0.005993 | 0.337401 |
| GSK3B | 7 | 0.011714 | 0.361817 |
| MET | 7 | 0.015481 | 0.373312 |
| ACHE | 7 | 0.01161 | 0.321697 |
| CA12 | 7 | 0.00754 | 0.340827 |
| ESR2 | 7 | 0.006213 | 0.322138 |
| ABCG2 | 7 | 0.005568 | 0.315648 |
| ADORA2A | 7 | 0.006582 | 0.333097 |
| HSP90AA1 | 6 | 0.018593 | 0.342815 |
| CA1 | 6 | 0.005914 | 0.339841 |
| ALOX15 | 6 | 0.006983 | 0.356331 |
| CA4 | 6 | 0.00177 | 0.312708 |
| CA7 | 6 | 0.00177 | 0.312708 |
| CA2 | 6 | 0.004935 | 0.332626 |
| TOP1 | 6 | 0.011607 | 0.355791 |
| PTGS2 | 6 | 0.006257 | 0.324362 |
| RORC | 6 | 0.008748 | 0.296905 |
| BACE1 | 6 | 0.005217 | 0.341322 |
| HSD17B1 | 6 | 0.001522 | 0.310641 |
| NOX4 | 6 | 0.004569 | 0.334044 |
| AKR1B1 | 6 | 0.011254 | 0.312292 |
| AKR1B10 | 6 | 0.010583 | 0.345843 |
| CA9 | 5 | 0.00332 | 0.334044 |
| ESR1 | 5 | 0.004809 | 0.313542 |
| TOP2A | 5 | 0.005732 | 0.315648 |
| PLK1 | 5 | 0.005426 | 0.353118 |
| CA14 | 5 | 0.004607 | 0.339841 |
| PIM1 | 5 | 0.005426 | 0.353118 |
| AURKB | 5 | 0.005426 | 0.353118 |
| SYK | 5 | 0.005994 | 0.345334 |
| KDR | 5 | 0.006196 | 0.347376 |
| XDH | 5 | 0.000736 | 0.298792 |
| SLC6A2 | 5 | 0.004667 | 0.299173 |
| SIGMAR1 | 5 | 0.02124 | 0.344322 |
| TERT | 5 | 0.006892 | 0.32481 |
| PLA2G1B | 5 | 0.004641 | 0.313542 |
| APP | 5 | 0.002584 | 0.305393 |
| ADORA3 | 5 | 0.003464 | 0.295412 |
| FLT3 | 5 | 0.003596 | 0.332626 |
| ADORA1 | 5 | 0.001511 | 0.298792 |
| PTPN1 | 5 | 0.004758 | 0.300319 |
| HSD11B1 | 5 | 0.008059 | 0.329362 |
| LGALS8 | 4 | 0.000193 | 0.266289 |
| LGALS4 | 4 | 0.000193 | 0.266289 |
| HTR6 | 4 | 0.000193 | 0.266289 |
| DRD1 | 4 | 0.000193 | 0.266289 |
| HTR2B | 4 | 0.000193 | 0.266289 |
| HPSE | 4 | 0.000193 | 0.266289 |
| FGF2 | 4 | 0.000193 | 0.266289 |
| FGF1 | 4 | 0.000193 | 0.266289 |
| VEGFA | 4 | 0.000193 | 0.266289 |
| PSEN2 | 4 | 0.000193 | 0.266289 |
| AKT1 | 4 | 0.003175 | 0.317782 |
| SRC | 4 | 0.002169 | 0.326162 |
| CA5A | 4 | 0.001048 | 0.309007 |
| CA6 | 4 | 0.001048 | 0.309007 |
| PTPRS | 4 | 0.000408 | 0.295784 |
| HTR2C | 4 | 0.002261 | 0.282622 |
| MAOA | 4 | 0.001048 | 0.309007 |
| F2 | 4 | 0.002809 | 0.302251 |
| PTK2 | 4 | 0.003218 | 0.321697 |
| RAF1 | 4 | 0.003688 | 0.336918 |
| CDK2 | 4 | 0.003841 | 0.342316 |
| ALPL | 4 | 0.00617 | 0.337401 |
| MAOB | 4 | 0.002862 | 0.32526 |
| TNKS2 | 4 | 0.001498 | 0.300704 |
| ABCC1 | 4 | 0.000912 | 0.304601 |
| CHRM2 | 4 | 0.002093 | 0.283645 |
| PPARA | 4 | 0.001619 | 0.293567 |
| CYP1B1 | 4 | 0.000358 | 0.291383 |
| ADRA2C | 3 | 0.002826 | 0.26965 |
| CYP2D6 | 3 | 7.28E-05 | 0.263897 |
| ADRA2B | 3 | 7.28E-05 | 0.263897 |
| PARP1 | 3 | 0.00502 | 0.295412 |
| MMP13 | 3 | 0.000502 | 0.301475 |
| PIK3R1 | 3 | 0.001225 | 0.315225 |
| CDK5R1 | 3 | 0.000502 | 0.301475 |
| GLO1 | 3 | 0.000502 | 0.301475 |
| CA13 | 3 | 0.000292 | 0.295412 |
| CA3 | 3 | 0.000292 | 0.295412 |
| TNKS | 3 | 0.000292 | 0.295412 |
| TLR9 | 3 | 0.000349 | 0.287814 |
| MIF | 3 | 0.000789 | 0.299554 |
| ESRRA | 3 | 0.000292 | 0.295412 |
| AVPR2 | 3 | 0.000639 | 0.289587 |
| ADRA1D | 3 | 0.001421 | 0.267806 |
| AURKA | 3 | 0.001804 | 0.323024 |
| MMP8 | 3 | 0.001804 | 0.323024 |
| MKNK1 | 3 | 0.004177 | 0.322138 |
| CHEK1 | 3 | 0.001804 | 0.323024 |
| TYMS | 3 | 0.00693 | 0.298034 |
| WEE1 | 3 | 0.001804 | 0.323024 |
| ABL1 | 3 | 0.001804 | 0.323024 |
| PIK3CA | 3 | 0.000645 | 0.289231 |
| CASP7 | 3 | 0.002906 | 0.302251 |
| IGF1R | 3 | 0.000219 | 0.289944 |
| AGTR1 | 3 | 0.001754 | 0.297657 |
| NR1H3 | 3 | 0.000216 | 0.280932 |
| SLC6A4 | 3 | 0.001009 | 0.280932 |
| PGR | 3 | 0.001009 | 0.280932 |
| SLC6A3 | 3 | 0.001009 | 0.280932 |
| HMGCR | 3 | 0.000216 | 0.280932 |
| NR3C1 | 3 | 0.001009 | 0.280932 |
| CYP17A1 | 3 | 0.000216 | 0.280932 |
| NPC1L1 | 3 | 0.000216 | 0.280932 |
| CYP51A1 | 3 | 0.000216 | 0.280932 |
| AR | 3 | 0.000216 | 0.280932 |
| NR3C2 | 3 | 0.001009 | 0.280932 |
| CDC25A | 3 | 0.004224 | 0.310231 |
| CES2 | 3 | 0.00154 | 0.287814 |
| NOS2 | 3 | 0.001871 | 0.281269 |
| PTGES | 3 | 0.001754 | 0.297657 |
| PDE4D | 3 | 0.00154 | 0.287814 |
| CDC25B | 3 | 0.004224 | 0.310231 |
| OPRD1 | 3 | 0.00309 | 0.274052 |
| SLC22A12 | 3 | 0.000112 | 0.288521 |
| POLB | 3 | 0.00025 | 0.276308 |
| GLI1 | 2 | 0.0004 | 0.275337 |
| GLI2 | 2 | 0.0004 | 0.275337 |
| TRPV1 | 2 | 0.000805 | 0.263601 |
| HTR1B | 2 | 1.14E-05 | 0.260966 |
| LGALS3 | 2 | 1.14E-05 | 0.260966 |
| STAT3 | 2 | 3.57E-05 | 0.263305 |
| SF3B3 | 2 | 3.57E-05 | 0.263305 |
| TRPV4 | 2 | 2.99E-05 | 0.262131 |
| MLNR | 2 | 2.99E-05 | 0.262131 |
| PPM1A | 2 | 2.99E-05 | 0.262131 |
| PTAFR | 2 | 2.99E-05 | 0.262131 |
| ADRA2A | 2 | 3.04E-05 | 0.261547 |
| GLRA2 | 2 | 2.99E-05 | 0.262131 |
| GLRA1 | 2 | 2.99E-05 | 0.262131 |
| CTSL | 2 | 0.000907 | 0.28676 |
| BCHE | 2 | 0.00154 | 0.287814 |
| SLC5A1 | 2 | 0.000319 | 0.262423 |
| CCNE1 | 2 | 0 | 0.283988 |
| BRAF | 2 | 0.000447 | 0.299554 |
| PDK1 | 2 | 0.000447 | 0.299554 |
| PDE7A | 2 | 0.001386 | 0.29653 |
| CASP3 | 2 | 0.001386 | 0.29653 |
| DNM1 | 2 | 0.000447 | 0.299554 |
| PIK3CG | 2 | 0.000607 | 0.312292 |
| PIK3CB | 2 | 0.000766 | 0.28676 |
| PIK3CD | 2 | 0.000766 | 0.28676 |
| MTOR | 2 | 0.000447 | 0.299554 |
| BMP1 | 2 | 0.000447 | 0.299554 |
| EPHX2 | 2 | 0.000316 | 0.255574 |
| MAPT | 2 | 3.66E-06 | 0.287111 |
| CD38 | 2 | 3.66E-06 | 0.287111 |
| MMP12 | 2 | 3.66E-06 | 0.287111 |
| AKR1A1 | 2 | 3.66E-06 | 0.287111 |
| AKR1C4 | 2 | 3.66E-06 | 0.287111 |
| AKR1C3 | 2 | 3.66E-06 | 0.287111 |
| AKR1C1 | 2 | 3.66E-06 | 0.287111 |
| AKR1C2 | 2 | 3.66E-06 | 0.287111 |
| NUAK1 | 2 | 3.66E-06 | 0.287111 |
| AXL | 2 | 3.66E-06 | 0.287111 |
| NEK6 | 2 | 3.66E-06 | 0.287111 |
| ALK | 2 | 3.66E-06 | 0.287111 |
| CAMK2B | 2 | 3.66E-06 | 0.287111 |
| CXCR1 | 2 | 3.66E-06 | 0.287111 |
| NEK2 | 2 | 3.66E-06 | 0.287111 |
| PKN1 | 2 | 3.66E-06 | 0.287111 |
| MMP3 | 2 | 3.66E-06 | 0.287111 |
| PYGL | 2 | 3.66E-06 | 0.287111 |
| MPO | 2 | 3.66E-06 | 0.287111 |
| DRD4 | 2 | 3.66E-06 | 0.287111 |
| CSNK2A1 | 2 | 3.66E-06 | 0.287111 |
| CDK6 | 2 | 3.66E-06 | 0.287111 |
| TTR | 2 | 3.66E-06 | 0.287111 |
| MPG | 2 | 3.66E-06 | 0.287111 |
| DAPK1 | 2 | 3.66E-06 | 0.287111 |
| GPR35 | 2 | 3.66E-06 | 0.287111 |
| ARG1 | 2 | 3.66E-06 | 0.287111 |
| CCNB3 | 2 | 3.66E-06 | 0.287111 |
| MMP2 | 2 | 3.66E-06 | 0.287111 |
| MMP9 | 2 | 3.66E-06 | 0.287111 |
| AHR | 2 | 3.66E-06 | 0.287111 |
| CA5B | 2 | 0.000258 | 0.287462 |
| PFKFB3 | 2 | 0.000139 | 0.292835 |
| CBR1 | 2 | 8.41E-06 | 0.270581 |
| HTR2A | 2 | 0.000693 | 0.277942 |
| ALDH2 | 2 | 8.41E-06 | 0.270581 |
| CHRM4 | 2 | 0.001674 | 0.265988 |
| PRSS1 | 2 | 6.19E-05 | 0.258384 |
| RXRA | 2 | 6.19E-05 | 0.258384 |
| CHRM1 | 2 | 0.000362 | 0.261838 |
| CHRNA7 | 2 | 0.003492 | 0.291022 |
| DRD2 | 2 | 0.001663 | 0.292835 |
| TAAR1 | 2 | 0.000717 | 0.308197 |
| DYRK1B | 2 | 0.002073 | 0.298413 |
| CLK3 | 2 | 0.00064 | 0.301863 |
| EPHB2 | 2 | 0.000717 | 0.308197 |
| LCK | 2 | 0.000717 | 0.308197 |
| RPS6KB1 | 2 | 0.00064 | 0.301863 |
| CDK4 | 2 | 0.000717 | 0.308197 |
| JAK3 | 2 | 0.00068 | 0.287814 |
| MMP16 | 2 | 0.000717 | 0.308197 |
| PIM2 | 2 | 0.000717 | 0.308197 |
| PDE10A | 2 | 0.000717 | 0.308197 |
| MMP7 | 2 | 0.000717 | 0.308197 |
| RET | 2 | 0.000717 | 0.308197 |
| ADAM17 | 2 | 0.000717 | 0.308197 |
| MMP1 | 2 | 0.000717 | 0.308197 |
| F3 | 2 | 0.00064 | 0.301863 |
| ALOX15B | 2 | 0.000717 | 0.308197 |
| PDGFRA | 2 | 0.000847 | 0.290662 |
| CMA1 | 2 | 0.000847 | 0.290662 |
| FGFR1 | 2 | 0.000847 | 0.290662 |
| ALOX5AP | 2 | 0 | 0.27566 |
| RORA | 2 | 0 | 0.27566 |
| GLUL | 2 | 0 | 0.27566 |
| IL6 | 2 | 0 | 0.27566 |
| PTGIR | 2 | 0 | 0.27566 |
| NR1H4 | 2 | 0 | 0.27566 |
| PTGDR2 | 2 | 0 | 0.27566 |
| GPBAR1 | 2 | 0 | 0.27566 |
| GRIK2 | 2 | 0 | 0.27566 |
| GRIK1 | 2 | 0 | 0.27566 |
| AMPD2 | 2 | 0 | 0.27566 |
| PTGER4 | 2 | 0 | 0.27566 |
| PRKCH | 2 | 0 | 0.27566 |
| PTGER2 | 2 | 0 | 0.27566 |
| PTPN11 | 2 | 0 | 0.27566 |
| FABP5 | 2 | 0 | 0.27566 |
| FABP3 | 2 | 0 | 0.27566 |
| FABP4 | 2 | 0 | 0.27566 |
| G6PD | 2 | 0 | 0.27566 |
| SHBG | 2 | 0 | 0.27566 |
| SERPINA6 | 2 | 0 | 0.27566 |
| HSD11B2 | 2 | 0 | 0.27566 |
| PPARD | 2 | 0 | 0.27566 |
| FAAH | 2 | 0 | 0.27566 |
| FNTA | 2 | 0 | 0.27566 |
| LTB4R | 2 | 0 | 0.27566 |
| PPARG | 2 | 0 | 0.27566 |
| PREP | 2 | 0 | 0.27566 |
| SCD | 2 | 0 | 0.27566 |
| FABP1 | 2 | 0 | 0.27566 |
| PTPN6 | 2 | 0 | 0.27566 |
| ACP1 | 2 | 0 | 0.27566 |
| PTPN2 | 2 | 0 | 0.27566 |
| PTPRF | 2 | 0 | 0.27566 |
| CD81 | 2 | 0 | 0.27566 |
| MCL1 | 2 | 9.49E-05 | 0.277942 |
| P2RX7 | 1 | 0 | 0.242894 |
| HCRTR1 | 1 | 0 | 0.242894 |
| HCRTR2 | 1 | 0 | 0.242894 |
| KCNA5 | 1 | 0 | 0.242894 |
| IDO1 | 1 | 0 | 0.242894 |
| PTK2B | 1 | 0 | 0.242894 |
| TRPV3 | 1 | 0 | 0.242894 |
| OPRK1 | 1 | 0 | 0.242894 |
| OPRM1 | 1 | 0 | 0.242894 |
| MDM2 | 1 | 0 | 0.242894 |
| HTR7 | 1 | 0 | 0.241149 |
| ADH1A | 1 | 0 | 0.241149 |
| STAT6 | 1 | 0 | 0.272464 |
| DPP4 | 1 | 0 | 0.272464 |
| ROCK2 | 1 | 0 | 0.272464 |
| CHEK2 | 1 | 0 | 0.272464 |
| PDGFRB | 1 | 0 | 0.272464 |
| NFE2L2 | 1 | 0 | 0.272464 |
| SERPINE1 | 1 | 0 | 0.272464 |
| EEF2K | 1 | 0 | 0.272464 |
| PLCG1 | 1 | 0 | 0.272464 |
| IKBKG | 1 | 0 | 0.272464 |
| EP300 | 1 | 0 | 0.272464 |
| CYP2C19 | 1 | 0 | 0.234883 |
| CYP3A4 | 1 | 0 | 0.234883 |
| PRKDC | 1 | 0 | 0.234883 |
| ERN1 | 1 | 0 | 0.234883 |
| ADRA1A | 1 | 0 | 0.258668 |
| DRD3 | 1 | 0 | 0.258668 |
| CNR2 | 1 | 0 | 0.259812 |
| CNR1 | 1 | 0 | 0.259812 |
| TACR2 | 1 | 0 | 0.259812 |
| CAPN1 | 1 | 0 | 0.259812 |
| REN | 1 | 0 | 0.259812 |
| LGALS9 | 1 | 0 | 0.251471 |
| FDFT1 | 1 | 0 | 0.251471 |
| CASP6 | 1 | 0 | 0.263011 |
| CASP9 | 1 | 0 | 0.263011 |
| CASP4 | 1 | 0 | 0.263011 |
| DYRK2 | 1 | 0 | 0.263011 |
| CTSV | 1 | 0 | 0.263011 |
| DYRK1A | 1 | 0 | 0.263011 |
| MERTK | 1 | 0 | 0.263011 |
| PLA2G10 | 1 | 0 | 0.263011 |
| MGLL | 1 | 0 | 0.263011 |
| GSTM2 | 1 | 0 | 0.263011 |
| GSTP1 | 1 | 0 | 0.263011 |
| PLAU | 1 | 0 | 0.263011 |
| ELANE | 1 | 0 | 0.263011 |
| CES1 | 1 | 0 | 0.263011 |
| NQO2 | 1 | 0 | 0.263011 |
| ERBB2 | 1 | 0 | 0.263011 |
| DCTPP1 | 1 | 0 | 0.263011 |
| TGM2 | 1 | 0 | 0.263011 |
| PTPRC | 1 | 0 | 0.263011 |
| PDE7B | 1 | 0 | 0.263011 |
| PDE4B | 1 | 0 | 0.263011 |
| APEX1 | 1 | 0 | 0.284676 |
| MYLK | 1 | 0 | 0.284676 |
| INSR | 1 | 0 | 0.284676 |
| KDM4E | 1 | 0 | 0.284676 |
| HRAS | 1 | 0 | 0.251203 |
| SLC5A2 | 1 | 0 | 0.251203 |
| ECE1 | 1 | 0 | 0.283988 |
| MAPK1 | 1 | 0 | 0.283988 |
| ROCK1 | 1 | 0 | 0.283988 |
| RPS6KA1 | 1 | 0 | 0.283988 |
| CSNK1G1 | 1 | 0 | 0.283988 |
| DBF4 | 1 | 0 | 0.283988 |
| MBD2 | 1 | 0 | 0.283988 |
| CCNE2 | 1 | 0 | 0.283988 |
| CCND1 | 1 | 0 | 0.283988 |
| TBK1 | 1 | 0 | 0.283988 |
| PIM3 | 1 | 0 | 0.283988 |
| COMT | 1 | 0 | 0.283988 |
| LNPEP | 1 | 0 | 0.283988 |
| MAPKAPK2 | 1 | 0 | 0.283988 |
| ADAM10 | 1 | 0 | 0.283988 |
| MMP25 | 1 | 0 | 0.283988 |
| FLT1 | 1 | 0 | 0.283988 |
| VCP | 1 | 0 | 0.283988 |
| HDAC10 | 1 | 0 | 0.283988 |
| HDAC9 | 1 | 0 | 0.283988 |
| HDAC4 | 1 | 0 | 0.283988 |
| HDAC11 | 1 | 0 | 0.283988 |
| NCOR1 | 1 | 0 | 0.283988 |
| CTSS | 1 | 0 | 0.283988 |
| HDAC7 | 1 | 0 | 0.283988 |
| HDAC5 | 1 | 0 | 0.283988 |
| NCOR2 | 1 | 0 | 0.283988 |
| CCND3 | 1 | 0 | 0.283988 |
| HDAC2 | 1 | 0 | 0.283988 |
| HDAC3 | 1 | 0 | 0.283988 |
| HSD17B3 | 1 | 0 | 0.283988 |
| RPS6KA2 | 1 | 0 | 0.283988 |
| NR4A1 | 1 | 0 | 0.283988 |
| HTT | 1 | 0 | 0.283988 |
| EZR | 1 | 0 | 0.283988 |
| PYGM | 1 | 0 | 0.253096 |
| LDHA | 1 | 0 | 0.253096 |
| ATIC | 1 | 0 | 0.253096 |
| GART | 1 | 0 | 0.253096 |
| FPGS | 1 | 0 | 0.253096 |
| HDAC1 | 1 | 0 | 0.253096 |
| HDAC8 | 1 | 0 | 0.253096 |
| SLC46A1 | 1 | 0 | 0.253096 |
| HDAC6 | 1 | 0 | 0.253096 |
| FOLR2 | 1 | 0 | 0.253096 |
| FOLR1 | 1 | 0 | 0.253096 |
| SLC19A1 | 1 | 0 | 0.253096 |
| DHFR | 1 | 0 | 0.253096 |
| GRK6 | 1 | 0 | 0.284676 |
| AMY1A | 1 | 0 | 0.284676 |
| CFTR | 1 | 0 | 0.284676 |
| FEN1 | 1 | 0 | 0.27027 |
| ERCC5 | 1 | 0 | 0.27027 |
| STS | 1 | 0 | 0.27027 |
| DHODH | 1 | 0 | 0.27027 |
| PON1 | 1 | 0 | 0.27027 |
| ESRRB | 1 | 0 | 0.27027 |
| MGAM | 1 | 0 | 0.27027 |
| TBXAS1 | 1 | 0 | 0.27027 |
| IL2 | 1 | 0 | 0.27027 |
| JUN | 1 | 0 | 0.252281 |
| PDE8B | 1 | 0 | 0.252281 |
| HSP90AB1 | 1 | 0 | 0.252281 |
| AVPR1A | 1 | 0 | 0.252281 |
| OXTR | 1 | 0 | 0.252281 |
| PNRC1 | 1 | 0 | 0.257534 |
| PNRC2 | 1 | 0 | 0.257534 |
| GABRA1 | 1 | 0 | 0.257534 |
| ADRB2 | 1 | 0 | 0.257534 |
| ADRA1B | 1 | 0 | 0.257534 |
| HTR3A | 1 | 0 | 0.257534 |
| SCN5A | 1 | 0 | 0.257534 |
| CHRM3 | 1 | 0 | 0.257534 |
| Ncoa1 | 1 | 0 | 0.250934 |
| CDC25C | 1 | 0 | 0.284676 |
| PDF | 1 | 0 | 0.284676 |
| FKBP1A | 1 | 0 | 0.284676 |
| EPHA1 | 1 | 0 | 0.284676 |
| COQ8B | 1 | 0 | 0.284676 |
| TYRO3 | 1 | 0 | 0.284676 |
| BTK | 1 | 0 | 0.284676 |
| EPHA3 | 1 | 0 | 0.284676 |
| EPHB3 | 1 | 0 | 0.284676 |
| PTK6 | 1 | 0 | 0.284676 |
| EPHA6 | 1 | 0 | 0.284676 |
| FGR | 1 | 0 | 0.284676 |
| TXK | 1 | 0 | 0.284676 |
| CLK2 | 1 | 0 | 0.284676 |
| CLK1 | 1 | 0 | 0.284676 |
| CLK4 | 1 | 0 | 0.284676 |
| FASN | 1 | 0 | 0.284676 |
| EPHA4 | 1 | 0 | 0.284676 |
| EPHA5 | 1 | 0 | 0.284676 |
| LYN | 1 | 0 | 0.284676 |
| BMX | 1 | 0 | 0.284676 |
| CSK | 1 | 0 | 0.284676 |
| BLK | 1 | 0 | 0.284676 |
| ATP4B | 1 | 0 | 0.284676 |
| YES1 | 1 | 0 | 0.284676 |
| EPHA2 | 1 | 0 | 0.284676 |
| FYN | 1 | 0 | 0.284676 |
| LDHB | 1 | 0 | 0.284676 |
| PDE2A | 1 | 0 | 0.284676 |
| TRPM8 | 1 | 0 | 0.284676 |
| IMPDH2 | 1 | 0 | 0.284676 |
| QPCT | 1 | 0 | 0.284676 |
| PPIA | 1 | 0 | 0.284676 |
| TGFBR1 | 1 | 0 | 0.284676 |
| GCGR | 1 | 0 | 0.284676 |
| EGLN1 | 1 | 0 | 0.284676 |
| ANPEP | 1 | 0 | 0.284676 |
| NDUFA4 | 1 | 0 | 0.284676 |
| FLT4 | 1 | 0 | 0.284676 |
| GUSB | 1 | 0 | 0.284676 |
| KIT | 1 | 0 | 0.284676 |
| NEK1 | 1 | 0 | 0.284676 |
| ADAMTS5 | 1 | 0 | 0.284676 |
| PLA2G7 | 1 | 0 | 0.284676 |
| SCN10A | 1 | 0 | 0.254743 |
| PDE3A | 1 | 0 | 0.254743 |
| BAZ2A | 1 | 0 | 0.254743 |
| TTK | 1 | 0 | 0.254743 |
| SIRT1 | 1 | 0 | 0.254743 |
| PLAT | 1 | 0 | 0.254743 |
| BAZ2B | 1 | 0 | 0.254743 |
| MAP2K1 | 1 | 0 | 0.254743 |
| SAE1 | 1 | 0 | 0.250133 |

**Table S3**. Gene of PPI network

| Name | Neighborhood Connectivity | Number Of Directed Edges | Degree |
| --- | --- | --- | --- |
| RORC | 4 | 1 | 1 |
| TNKS2 | 15 | 2 | 2 |
| HTR6 | 19 | 1 | 1 |
| NPC1L1 | 21.5 | 4 | 4 |
| CYP51A1 | 19.66666667 | 3 | 3 |
| HTR2C | 12.375 | 8 | 8 |
| HSD11B1 | 15.85714286 | 7 | 7 |
| POLB | 21.33333333 | 3 | 3 |
| TNKS | 17.33333333 | 6 | 6 |
| PIM1 | 28 | 4 | 4 |
| CA14 | 6.66666667 | 3 | 3 |
| CA12 | 16.75 | 4 | 4 |
| CA7 | 4.75 | 4 | 4 |
| CA13 | 4.75 | 4 | 4 |
| CA3 | 4.75 | 4 | 4 |
| CA2 | 6 | 5 | 5 |
| PLK1 | 22.76470588 | 17 | 17 |
| AURKB | 20.5 | 14 | 14 |
| TOP2A | 25.05882353 | 17 | 17 |
| HSD17B1 | 18.09090909 | 11 | 11 |
| CYP17A1 | 19.27272727 | 11 | 11 |
| STS | 21.57142857 | 7 | 7 |
| HSD17B2 | 20.55555556 | 9 | 9 |
| CA4 | 10 | 4 | 4 |
| ALPL | 7 | 1 | 1 |
| SYK | 22.2 | 10 | 10 |
| PTGES | 19.5 | 10 | 10 |
| ALOX12 | 14.71428571 | 7 | 7 |
| ALOX15 | 21.625 | 8 | 8 |
| FGF1 | 32.33333333 | 12 | 12 |
| NR1H3 | 25.75 | 4 | 4 |
| WEE1 | 28.46153846 | 13 | 13 |
| CHEK1 | 25.9047619 | 21 | 21 |
| ESRRA | 39.33333333 | 6 | 6 |
| CYP19A1 | 25.22727273 | 22 | 22 |
| CA9 | 28.08333333 | 12 | 12 |
| CASP7 | 36.11111111 | 9 | 9 |
| PTGS1 | 23.55555556 | 9 | 9 |
| TLR9 | 33.28571429 | 7 | 7 |
| PDE4D | 42.33333333 | 3 | 3 |
| ESR2 | 29.82352941 | 17 | 17 |
| PTK2 | 28.10526316 | 19 | 19 |
| NOS2 | 29.36363636 | 11 | 11 |
| CDK5R1 | 28.6 | 10 | 10 |
| PLA2G1B | 21.57142857 | 7 | 7 |
| CDC25A | 23.23529412 | 17 | 17 |
| LGALS4 | 37 | 9 | 9 |
| SIGMAR1 | 23.66666667 | 6 | 6 |
| MMP13 | 29.28571429 | 7 | 7 |
| CDC25B | 24.46666667 | 15 | 15 |
| MMP8 | 40.83333333 | 6 | 6 |
| AURKA | 28.77777778 | 18 | 18 |
| MIF | 50.4 | 5 | 5 |
| AKR1B10 | 7.5 | 2 | 2 |
| GLO1 | 9.5 | 4 | 4 |
| XDH | 7.33333333 | 3 | 3 |
| PPARA | 20.82608696 | 23 | 23 |
| NR3C2 | 22 | 8 | 8 |
| F2 | 23.75 | 12 | 12 |
| NOX4 | 30.88888889 | 9 | 9 |
| HTR2B | 12.5 | 6 | 6 |
| RAF1 | 28.5 | 20 | 20 |
| AGTR1 | 27.66666667 | 12 | 12 |
| ADRA2C | 14.66666667 | 3 | 3 |
| AVPR2 | 10 | 5 | 5 |
| ADRA2B | 7.5 | 2 | 2 |
| ADRA1D | 14.66666667 | 3 | 3 |
| SLC6A2 | 7.3 | 10 | 10 |
| DRD1 | 11.85714286 | 7 | 7 |
| ADORA3 | 33.4 | 5 | 5 |
| ADORA2A | 26.42857143 | 7 | 7 |
| ADORA1 | 25.75 | 8 | 8 |
| BACE1 | 27 | 9 | 9 |
| MAOB | 10.57142857 | 14 | 14 |
| MAOA | 12 | 11 | 11 |
| CA1 | 6.83333333 | 6 | 6 |
| CHRM2 | 12.75 | 4 | 4 |
| PSEN2 | 17 | 4 | 4 |
| GSK3B | 26.2 | 25 | 25 |
| SLC6A3 | 16.53333333 | 15 | 15 |
| TYR | 31.4 | 5 | 5 |
| ACHE | 18.38461538 | 13 | 13 |
| PIK3R1 | 27.83333333 | 24 | 24 |
| CDK1 | 25.15384615 | 26 | 26 |
| ALOX5 | 22.81818182 | 11 | 11 |
| PTPN1 | 29.1 | 20 | 20 |
| PARP1 | 25.54166667 | 24 | 24 |
| PTPRS | 36.66666667 | 3 | 3 |
| TERT | 30.64705882 | 17 | 17 |
| APP | 26.47619048 | 21 | 21 |
| CDK2 | 25.03846154 | 26 | 26 |
| PIK3CA | 26.21875 | 32 | 32 |
| FLT3 | 28.69230769 | 13 | 13 |
| SLC22A12 | 11 | 2 | 2 |
| FGF2 | 26.76 | 25 | 25 |
| MET | 29.61111111 | 18 | 18 |
| CES2 | 19 | 2 | 2 |
| IGF1R | 31.23809524 | 21 | 21 |
| PGR | 26.76923077 | 26 | 26 |
| AR | 24.6969697 | 33 | 33 |
| VEGFA | 20.38461538 | 52 | 52 |
| CYP1B1 | 25.5 | 10 | 10 |
| AKT1 | 17.53731343 | 67 | 67 |
| ESR1 | 21.64583333 | 48 | 48 |
| ABCC1 | 29 | 7 | 7 |
| SRC | 22.7804878 | 41 | 41 |
| ABL1 | 27.57894737 | 19 | 19 |
| PTGS2 | 21.02564103 | 39 | 39 |
| TOP1 | 25.47368421 | 19 | 19 |
| CYP2D6 | 16.66666667 | 12 | 12 |
| HSP90AA1 | 22.15909091 | 44 | 44 |
| TYMS | 25.625 | 16 | 16 |
| HMGCR | 17.77777778 | 9 | 9 |
| AKR1B1 | 26 | 11 | 11 |
| EGFR | 20.48 | 50 | 50 |
| KDR | 28.47619048 | 21 | 21 |
| SLC6A4 | 10.94736842 | 19 | 19 |
| ABCG2 | 25.78947368 | 19 | 19 |
| OPRD1 | 15.5 | 4 | 4 |
| NR3C1 | 26.29166667 | 24 | 24 |
| ABCB1 | 26.55 | 20 | 20 |

**Table S4**. Total score and composition of pathway enrichment in 8 subcategories

| Network | Symbol | Total Score | Number of symbols |
| --- | --- | --- | --- |
| MCODE 1 | PLK1 | 66 | 20 |
|  | PTPN1 |  |  |
|  | ABL1 |  |  |
|  | ADRA2C |  |  |
|  | SYK |  |  |
|  | WEE1 |  |  |
|  | ADRA2B |  |  |
|  | CDC25A |  |  |
|  | CHRM2 |  |  |
|  | CDC25B |  |  |
|  | ADORA1 |  |  |
|  | CHEK1 |  |  |
|  | NR3C1 |  |  |
|  | APP |  |  |
|  | CDK2 |  |  |
|  | OPRD1 |  |  |
|  | EGFR |  |  |
|  | RAF1 |  |  |
|  | AKT1 |  |  |
|  | ADORA3 |  |  |
| MCODE 2 | IGF1R | 65.8 | 14 |
|  | PIK3CA |  |  |
|  | FGF1 |  |  |
|  | PIK3R1 |  |  |
|  | FGF2 |  |  |
|  | HSP90AA1 |  |  |
|  | ESR2 |  |  |
|  | ESR1 |  |  |
|  | MET |  |  |
|  | KDR |  |  |
|  | SRC |  |  |
|  | VEGFA |  |  |
|  | PTK2 |  |  |
|  | AR |  |  |
| MCODE 3 | F2 | 26 | 11 |
|  | PLA2G1B |  |  |
|  | ALOX5 |  |  |
|  | HTR2B |  |  |
|  | PTGS1 |  |  |
|  | ALOX15 |  |  |
|  | AGTR1 |  |  |
|  | ALOX12 |  |  |
|  | PTGS2 |  |  |
|  | HTR2C |  |  |
|  | ADRA1D |  |  |
| MCODE 4 | CYP2D6 | 7 | 6 |
|  | CYP51A1 |  |  |
|  | HSD17B1 |  |  |
|  | SIGMAR1 |  |  |
|  | CYP1B1 |  |  |
|  | HSD17B2 |  |  |
| MCODE 5 | CA2 | 10 | 5 |
|  | CA3 |  |  |
|  | CA7 |  |  |
|  | CA1 |  |  |
|  | CA13 |  |  |
| MCODE 6 | HTR6 | 6 | 4 |
|  | DRD1 |  |  |
|  | AVPR2 |  |  |
|  | ADORA2A |  |  |
| MCODE 7 | CA4 | 3 | 3 |
|  | CA14 |  |  |
|  | CA12 |  |  |
| MCODE 8 | PARP1 | 3 | 3 |
|  | TOP1 |  |  |
|  | TOP2A |  |  |

**Table S5**. The top 3 KEGG Pathways of 8 Subclasses Classified by MCODE Algorithm

| Network | KEGG | Annotation |
| --- | --- | --- |
| MCODE 1 | hsa04110 | Cell cycle |
|  | hsa04022 | cGMP-PKG signaling pathway |
|  | hsa04914 | Progesterone-mediated oocyte maturation |
| MCODE 2 | hsa05200 | Pathways in cancer |
|  | hsa05205 | Proteoglycans in cancer |
|  | hsa01521 | EGFR tyrosine kinase inhibitor resistance |
| MCODE 3 | hsa04726 | Serotonergic synapse |
|  | hsa00590 | Arachidonic acid metabolism |
|  | hsa04080 | Neuroactive ligand-receptor interaction |
| MCODE 4 | hsa04913 | Ovarian steroidogenesis |
|  | hsa00140 | Steroid hormone biosynthesis |
| MCODE 5 | hsa00910 | Nitrogen metabolism |
| MCODE 6 | hsa04080 | Neuroactive ligand-receptor interaction |
|  | hsa04024 | cAMP signaling pathway |
|  | hsa04020 | Calcium signaling pathway |
| MCODE 7 | hsa00910 | Nitrogen metabolism |
| MCODE 8 | none | None |

**Table S6** Genes of CGS_ARCR_ and their protein names

| Gene | Entry | Protein names |
| --- | --- | --- |
| CHEK1 | O14757 | Serine/threonine-protein kinase Chk1 |
| ABL1 | P00519 | Tyrosine-protein kinase ABL1 |
| EGFR | P00533 | Epidermal growth factor receptor |
| ESR1 | P03372 | Estrogen receptor |
| RAF1 | P04049 | RAF proto-oncogene serine/threonine-protein kinase |
| NR3C1 | P04150 | Glucocorticoid receptor |
| APP | P05067 | Amyloid-beta precursor protein |
| CDK1 | P06493 | Cyclin-dependent kinase 1 |
| HSP90AA1 | P07900 | Heat shock protein HSP 90-alpha |
| CHRM2 | P08172 | Muscarinic acetylcholine receptor M2 |
| ADORA3 | P0DMS8 | Adenosine receptor A3 |
| AR | P10275 | Androgen receptor |
| SRC | P12931 | Proto-oncogene tyrosine-protein kinase Src |
| VEGFA | P15692 | Vascular endothelial growth factor A, long form |
| PTPN1 | P18031 | Tyrosine-protein phosphatase non-receptor type 1 |
| ADRA2B | P18089 | Alpha-2B adrenergic receptor |
| ADRA2C | P18825 | Alpha-2C adrenergic receptor |
| CDK2 | P24941 | Cyclin-dependent kinase 2 |
| WEE1 | P30291 | Wee1-like protein kinase |
| CDC25A | P30304 | M-phase inducer phosphatase 1 |
| CDC25B | P30305 | M-phase inducer phosphatase 2 |
| ADORA1 | P30542 | Adenosine receptor A1 |
| AKT1 | P31749 | RAC-alpha serine/threonine-protein kinase |
| PTGS2 | P35354 | Prostaglandin G/H synthase 2 |
| OPRD1 | P41143 | Delta-type opioid receptor |
| PIK3CA | P42336 | Phosphatidylinositol 4,5-bisphosphate 3-kinase catalytic subunit alpha isoform |
| SYK | P43405 | Tyrosine-protein kinase SYK |
| PLK1 | P53350 | Serine/threonine-protein kinase PLK1 |

**Table S7** Genes of **CGS**ARCR and Corresponding Components of ARCR

| Ingredient | ID | Gene |
| --- | --- | --- |
| Formononetin | HQ9 | PTPN1 |
|  |  | ADORA1 |
|  |  | EGFR |
|  |  | RAF1 |
|  |  | ESR1 |
| Bifendate | HQ8 | SYK |
|  |  | EGFR |
| (6aR,11aR)-9,10-dimethoxy-6a,11a-dihydro-6H-benzofurano[3,2-c] chromen-3-ol | HQ7 | ESR1 |
| 9,10-dimethoxypterocarpan-3-O-β-D-glucoside | HQ6 | PTGS2 |
| 7-O-methylisomucronulatol | HQ5 | PLK1 |
|  |  | ABL1 |
|  |  | SYK |
|  |  | WEE1 |
|  |  | CDC25A |
|  |  | CDC25B |
|  |  | CHEK1 |
|  |  | CDK2 |
|  |  | EGFR |
|  |  | RAF1 |
|  |  | HSP90AA1 |
|  |  | CDK1 |
| 3,9-di-O-methylnissolin | HQ4 | PTPN1 |
|  |  | PIK3CA |
| (3S,8S,9S,10R,13R,14S,17R)-10,13-dimethyl-17-[(2R,5S)-5-propan-2-yloctan-2-yl]-2,3,4,7,8,9,11,12,14,15,16,17-dodecahydro-1H-cyclopenta[a]phenanthren-3-ol | HQ3 | EGFR |
|  |  | AR |
| Astragaloside IV | HQ22 | ADRA2C |
|  |  | ADRA2B |
|  |  | HSP90AA1 |
|  |  | CDK1 |
|  |  | VEGFA |
| Astragaloside III | HQ21 | ADRA2C |
|  |  | ADRA2B |
|  |  | HSP90AA1 |
|  |  | CDK1 |
|  |  | VEGFA |
| Astragaloside II | HQ20 | OPRD1 |
|  |  | HSP90AA1 |
|  |  | CDK1 |
|  |  | VEGFA |
| Jaranol | HQ2 | ADORA1 |
|  |  | APP |
|  |  | OPRD1 |
|  |  | ADORA3 |
| Astragaloside I | HQ19 | ADRA2B |
|  |  | VEGFA |
| Astragalus polysaccharides | HQ18 | EGFR |
|  |  | ADORA3 |
|  |  | PTGS2 |
| Quercetin | HQ17 | PLK1 |
|  |  | SYK |
|  |  | ADORA1 |
|  |  | APP |
|  |  | CDK2 |
|  |  | EGFR |
|  |  | AKT1 |
|  |  | SRC |
|  |  | CDK1 |
| 1,7-Dihydroxy-3,9-dimethoxy pterocarpene | HQ16 | PTGS2 |
| isomucronulatol-7,2’-di-O-glucosiole | HQ15 | ADORA1 |
|  |  | ADORA3 |
| (3R)-3-(2-hydroxy-3,4-dimethoxyphenyl)chroman-7-ol | HQ14 | PLK1 |
|  |  | ABL1 |
|  |  | SYK |
|  |  | WEE1 |
|  |  | CHEK1 |
|  |  | CDK2 |
|  |  | EGFR |
|  |  | RAF1 |
|  |  | AKT1 |
|  |  | SRC |
|  |  | PIK3CA |
|  |  | PIK3CA |
|  |  | HSP90AA1 |
|  |  | CDK1 |
| Kaempferol | HQ12 | PLK1 |
|  |  | SYK |
|  |  | ADORA1 |
|  |  | APP |
|  |  | CDK2 |
|  |  | EGFR |
|  |  | AKT1 |
|  |  | ESR1 |
|  |  | SRC |
|  |  | PTGS2 |
|  |  | CDK1 |
| Calycosin | HQ11 | EGFR |
|  |  | ESR1 |
| Mairin | HQ1 | PTPN1 |
| Curcumol | ES4 | ADRA2C |
|  |  | CHRM2 |
|  |  | NR3C1 |
|  |  | OPRD1 |
| Germacron | ES3 | CHRM2 |
| Bisdemethoxycurcumin | ES2 | PLK1 |
|  |  | ABL1 |
|  |  | WEE1 |
|  |  | CHEK1 |
|  |  | APP |
|  |  | EGFR |
|  |  | RAF1 |
|  |  | ESR1 |
|  |  | SRC |
|  |  | HSP90AA1 |
|  |  | CDK1 |
| Wenjine | ES1 | APP |
|  |  | AKT1 |
| Hederagenin | A1 | PTPN1 |
|  |  | PTPN1 |
|  |  | CDC25A |
|  |  | CDC25A |
|  |  | CHRM2 |
|  |  | CHRM2 |
|  |  | CDC25B |
|  |  | CDC25B |
|  |  | NR3C1 |
|  |  | NR3C1 |
|  |  | ADORA3 |
|  |  | ADORA3 |
|  |  | PTGS2 |
|  |  | PTGS2 |
|  |  | AR |

**Table S8.** Differential GSVA

| **Cancer type** | **Tumor gsva** | **Normal gsva** | **log2FC** | **Pval** | **Diff gsva** |
| --- | --- | --- | --- | --- | --- |
| ESCA | 1.474806358 | 1.156410339 | 0.350872129 | 0.00007213 | 0.31839602 |
| HNSC | 1.523531161 | 1.249681565 | 0.285858484 | 2.84712E-12 | 0.273849596 |
| COAD | 2.045997761 | 1.798387845 | 0.186100376 | 4.59148E-15 | 0.247609915 |
| LUSC | 1.124498317 | 0.893366152 | 0.331958002 | 3.25227E-41 | 0.231132165 |
| STAD | 1.281224233 | 1.133128154 | 0.177211955 | 1.80233E-09 | 0.148096079 |
| THCA | 1.5885321 | 1.462341195 | 0.119414282 | 5.15411E-11 | 0.126190905 |
| LUAD | 1.085251323 | 0.979740011 | 0.147558316 | 5.2172E-19 | 0.105511311 |
| KIRC | 1.098307446 | 1.016794664 | 0.111253596 | 9.58191E-08 | 0.081512782 |
| BRCA | 0.947189356 | 0.902410624 | 0.069868816 | 4.14518E-07 | 0.044778732 |
| BLCA | 1.157701659 | 1.122911782 | 0.044018925 | 0.203646041 | 0.034789877 |
| KIRP | 1.079302933 | 1.04793615 | 0.042549033 | 0.117718179 | 0.031366783 |
| LIHC | 0.914777134 | 0.892172016 | 0.036098408 | 0.116785079 | 0.022605119 |
| KICH | 0.990490644 | 1.048500011 | -0.08211162 | 0.081803148 | -0.058009367 |
| PRAD | 0.871330122 | 0.987741729 | -0.18091444 | 4.73767E-09 | -0.116411606 |

**Table S9.** GSEA

| Cancer type | | ES | NES | Pval | Padj | Log2err |
| --- | --- | --- | --- | --- | --- | --- |
| LUAD | 0.484323316 | | 1.082392994 | 0.38534279 | 0.38534279 | 0.06568136 |
| LUSC | 0.472755529 | | 1.101742526 | 0.359748428 | 0.359748428 | 0.072179797 |
| STAD | 0.435967511 | | 1.056448821 | 0.388467375 | 0.388467375 | 0.077884011 |
| HNSC | 0.431350125 | | 1.091572173 | 0.345991561 | 0.345991561 | 0.103197467 |
| BRCA | 0.396935365 | | 0.961363549 | 0.52574103 | 0.52574103 | 0.064070375 |
| ESCA | 0.393418048 | | 0.952456514 | 0.552325581 | 0.552325581 | 0.058345278 |
| THCA | 0.340806702 | | 0.785523287 | 0.758725341 | 0.758725341 | 0.045679041 |
| COAD | 0.338056332 | | 0.861024357 | 0.671256454 | 0.671256454 | 0.057125851 |
| KIRC | 0.251313307 | | 0.60834642 | 0.953232462 | 0.953232462 | 0.030418051 |

**Table S10.** Results of univariate Cox regression analysis of **CGS**ACAR in THCA

| ID | p-value | HR | HR.95L | HR.95H |
| --- | --- | --- | --- | --- |
| PTPN1 | 0.000995414 | 39.019 | 4.403757 | 345.7235 |
| ADRA2C | 0.099283208 | 1.291129 | 0.952842 | 1.749519 |
| CDC25A | 0.12436412 | 0.543206 | 0.249419 | 1.183037 |
| CHRM2 | 0.002403424 | 8.636752 | 2.146425 | 34.75244 |
| CDC25B | 0.024328749 | 2.518414 | 1.127202 | 5.626686 |
| APP | 0.051085475 | 0.240759 | 0.057577 | 1.00674 |
| CDK2 | 0.057650517 | 0.172556 | 0.028125 | 1.058689 |
| RAF1 | 0.148927377 | 5.366231 | 0.548048 | 52.54368 |
| ADORA3 | 0.00023127 | 0.316191 | 0.171309 | 0.583606 |
| ESR1 | 0.026078837 | 1.586606 | 1.056531 | 2.382627 |
| PTGS2 | 0.007388634 | 0.568094 | 0.375612 | 0.859212 |
| PIK3CA | 0.103063966 | 3.351284 | 0.782978 | 14.34409 |
| HSP90AA1 | 0.046660802 | 0.137486 | 0.019465 | 0.971099 |

**Table S11.** Results of univariate cox regression analysis of CGS_ACAR_ in BRCA

| ID | p-value | HR | HR.95L | HR.95H |
| --- | --- | --- | --- | --- |
| PLK1 | 0.002742356 | 1.47 | 1.14 | 1.88 |
| ADRA2C | 0.012445591 | 1.12 | 1.02 | 1.22 |
| CDC25A | 0.144359873 | 0.83 | 0.65 | 1.07 |
| ADORA1 | 0.021398772 | 0.89 | 0.81 | 0.98 |
| OPRD1 | 0.118556423 | 1.10 | 0.98 | 1.23 |
| EGFR | 0.005792321 | 0.85 | 0.75 | 0.95 |
| ADORA3 | 0.004514425 | 1.28 | 1.08 | 1.52 |
| ESR1 | 0.121223703 | 0.94 | 0.88 | 1.02 |
| PIK3CA | 0.012133933 | 1.45 | 1.08 | 1.94 |
| HSP90AA1 | 0.109821925 | 1.23 | 0.95 | 1.59 |
| CDK1 | 0.121877308 | 0.82 | 0.63 | 1.06 |
| VEGFA | 0.081286223 | 1.19 | 0.98 | 1.45 |

**Table S12.** Results of univariate Cox regression analysis of **CGS**ACAR in LUAD

| ID | p-value | HR | HR.95L | HR.95H |
| --- | --- | --- | --- | --- |
| PLK1 | 2.62E-08 | 1.65919 | 1.388187 | 1.983098 |
| ABL1 | 0.019137903 | 1.446737 | 1.062212 | 1.970463 |
| ADRA2C | 0.022077353 | 1.100375 | 1.013846 | 1.194289 |
| SYK | 0.146902368 | 0.873212 | 0.727046 | 1.048763 |
| WEE1 | 0.096011292 | 1.210942 | 0.966597 | 1.517055 |
| CHRM2 | 0.111646337 | 0.873385 | 0.739212 | 1.031911 |
| CDC25B | 0.000334597 | 0.71915 | 0.600597 | 0.861105 |
| ADORA1 | 0.05452325 | 1.085328 | 0.998412 | 1.17981 |
| NR3C1 | 0.007366534 | 1.358106 | 1.085696 | 1.698867 |
| CDK2 | 0.15441223 | 0.789225 | 0.569799 | 1.09315 |

**Table S13.** Results of univariate Cox regression analysis of **CGS**_ACAR_ in KRIC

| ID | p-value | HR | HR.95L | HR.95H |
| --- | --- | --- | --- | --- |
| PLK1 | 1.06E-06 | 1.397797 | 1.221918 | 1.598993 |
| ADRA2C | 0.021757989 | 0.881824 | 0.792005 | 0.981828 |
| SYK | 0.059961858 | 0.862883 | 0.739976 | 1.006204 |
| APP | 0.129889881 | 0.795482 | 0.591611 | 1.069608 |
| ESR1 | 0.021047348 | 1.175893 | 1.024681 | 1.349418 |
| SRC | 0.00058274 | 1.626744 | 1.232826 | 2.146527 |
| PTGS2 | 0.001845293 | 1.16389 | 1.057858 | 1.280549 |
| HSP90AA1 | 0.014101484 | 0.702637 | 0.530092 | 0.931345 |
| AR | 0.000269061 | 0.847194 | 0.774892 | 0.926242 |

**Table S14.** Results of univariate Cox regression analysis of **CGS**ACAR in COAD

| ID | p-value | HR | HR.95L | HR.95H |
| --- | --- | --- | --- | --- |
| PLK1 | 0.026110524 | 0.596423 | 0.378273 | 0.94038 |
| ABL1 | 0.068973848 | 0.532184 | 0.269668 | 1.050256 |
| NR3C1 | 0.070330329 | 0.726321 | 0.513728 | 1.026891 |
| APP | 0.049901859 | 0.563986 | 0.318158 | 0.999755 |
| ESR1 | 0.016374506 | 1.453726 | 1.071066 | 1.973098 |
| PIK3CA | 0.063919906 | 1.910317 | 0.963233 | 3.788608 |
| VEGFA | 0.059010824 | 1.445984 | 0.986064 | 2.12042 |

**Table S15.** Results of univariate cox regression analysis of **CGS**ACAR in STAD

| ID | p-value | HR | HR.95L | HR.95H |
| --- | --- | --- | --- | --- |
| ADRA2C | 0.081910153 | 0.923924 | 0.845126 | 1.010068 |
| CHEK1 | 0.087792664 | 0.789337 | 0.601604 | 1.035652 |
| NR3C1 | 0.014781019 | 1.381956 | 1.065452 | 1.792481 |
| EGFR | 0.042900085 | 1.130038 | 1.003915 | 1.272007 |
| ESR1 | 0.108070891 | 0.875118 | 0.743714 | 1.029741 |
| AR | 0.038311628 | 1.151568 | 1.007628 | 1.31607 |
| CDK1 | 0.011179723 | 1.3668 | 1.073661 | 1.739974 |

**Table S16.** Results of univariate cox regression analysis of **CGS**ACAR in HNSC

| ID | p-value | HR | HR.95L | HR.95H |
| --- | --- | --- | --- | --- |
| PLK1 | 0.005312282 | 1.369193 | 1.09777 | 1.707726 |
| ADRA2C | 0.131259253 | 1.06024 | 0.982678 | 1.143922 |
| APP | 0.00017136 | 1.445646 | 1.192833 | 1.75204 |
| OPRD1 | 0.102233999 | 0.735791 | 0.509261 | 1.063088 |
| ESR1 | 0.020471335 | 0.905355 | 0.83234 | 0.984774 |
| PTGS2 | 0.027817033 | 0.930321 | 0.87234 | 0.992155 |
| HSP90AA1 | 0.151252923 | 1.218695 | 0.930223 | 1.596626 |

**Table S17.** Results of univariate cox regression analysis of **CGS**ACAR in ESCA

| ID | p-value | HR | HR.95L | HR.95H |
| --- | --- | --- | --- | --- |
| ADRA2B | 0.135750095 | 0.890577 | 0.764801 | 1.037037 |
| CDC25A | 0.043929923 | 1.505692 | 1.011193 | 2.242014 |
| CDC25B | 0.147376518 | 1.258151 | 0.922185 | 1.716516 |
| CHEK1 | 0.008517612 | 0.572978 | 0.378401 | 0.86761 |
| AKT1 | 0.002017832 | 0.399742 | 0.223355 | 0.715426 |
| HSP90AA1 | 0.038171382 | 1.596142 | 1.025827 | 2.483528 |

**Table S18**. Results of univariate cox regression analysis of **CGS**ACAR in LUSC

| ID | p-value | HR | HR.95L | HR.95H |
| --- | --- | --- | --- | --- |
| SYK | 0.115167508 | 1.136663 | 0.969213 | 1.333043 |
| WEE1 | 0.12822026 | 1.185961 | 0.951991 | 1.477434 |
| CDC25A | 0.079431511 | 0.852337 | 0.712975 | 1.018939 |
| CDC25B | 0.130764798 | 1.159791 | 0.956921 | 1.40567 |
| ADORA1 | 0.103405606 | 1.0941 | 0.981866 | 1.219162 |
| SRC | 0.094323235 | 1.222827 | 0.966086 | 1.547798 |

**Table S19.** Presence of 28 genes from the CGS_ARCR_ in seven potential indications

| Original id | BRCA-PS | COAD-PS | ESCA-PS | KRIC-PS | LUAD-PS | STAD-PS | THCA-PS |
| --- | --- | --- | --- | --- | --- | --- | --- |
| ADRA2C | √ | × | × | √ | √ | √ | √ |
| ESR1 | √ | √ | × | √ | × | √ | √ |
| PLK1 | √ | √ | × | √ | √ | × | × |
| HSP90AA1 | √ | × | √ | √ | × | × | √ |
| NR3C1 | × | √ | × | × | √ | √ | × |
| PIK3CA | √ | √ | × | × | × | × | √ |
| APP | × | √ | × | √ | × | × | √ |
| CDC25A | √ | × | √ | × | × | × | √ |
| CDC25B | × | × | √ | × | √ | × | √ |
| CDK1 | √ | × | × | × | × | √ | × |
| PTGS2 | × | × | × | √ | × | × | √ |
| CDK2 | × | × | × | × | √ | × | √ |
| ABL1 | × | √ | × | × | √ | × | × |
| CHRM2 | × | × | × | × | √ | × | √ |
| ADORA3 | √ | × | × | × | × | × | √ |
| SYK | × | × | × | √ | √ | × | × |
| ADORA1 | √ | × | × | × | √ | × | × |
| EGFR | √ | × | × | × | × | √ | × |
| CHEK1 | × | × | √ | × | × | √ | × |
| AR | × | × | × | √ | × | √ | × |
| VEGFA | √ | √ | × | × | × | × | × |
| PTPN1 | × | × | × | × | × | × | √ |
| WEE1 | × | × | × | × | √ | × | × |
| AKT1 | × | × | √ | × | × | × | × |
| ADRA2B | × | × | √ | × | × | × | × |
| RAF1 | × | × | × | × | × | × | √ |
| OPRD1 | √ | × | × | × | × | × | × |
| SRC | × | × | × | √ | × | × | × |

√: indicates that the gene exists in this gene list.

×: indicates that the gene does not exist in this gene list.

**Table S20.** Predictive Performance Ratings of Prognostic Models for Nine Potential Indications.

| **Cancer Type** | **AUC** | **Predicting Performance Reliability** |
| --- | --- | --- |
| THCA | > 0.85 for 1-7 years | High |
| LUAD | > 0.70 for 1-7 years | moderate |
| KRIC | > 0.70 for 1-7 years | moderate |
| COAD | > 0.70 for 4-7 years | general |
| BRCA | ≥ 0.70 for 3-5 and 7 years | general |
| STAD | ≥ 0.70 for 4-6 years | general |
| ESCA | > 0.70 for 3-5 consecutive years | general |
| HNSC | < 0.7 for 1-7 years | suboptimal |
| LUSC | < 0.7 for 1-7 years | suboptimal |

**Table S21.** Literature Reports on Prognostic Genes of ARCR in LUAD.

| **Gene** | **Feature** | **Summary** | **Citation** |
| --- | --- | --- | --- |
| PLK1 | ▲ | Wang *et al.* found that PLK1 was highly expressed in LUAD patients, and a potential prognostic factor associated with the tumor microenvironment in lung adenocarcinoma | PMID: 35941968 |
| ABL1 | ▲ | Gu *et al.* found that the expression of ABL1 kinases was significantly increased in three NSCLC tumor specimens compared to adjacent normal tissue from the same patient. | PMID: 28018973 |
| ADRA2C | - | No relevant reports were found. | - |
| SYK | ▼ | Duan *et al.* have discovered that the serum SYK levels of lung adenocarcinoma patients were remarkably lower than those of normal persons. | PMID: 20797333 |
| WEE1 | ▼ | Yoshida *et al.* discovered that the loss of Wee1 expression might have a potential role in promoting tumor progression and was a significant prognostic indicator in NSCLC by suppressing the activity of the Cyclin B1/cdc2 complex. | PMID: 14760118 |
| CHRM2 | - | No relevant reports were found. | - |
| CDC25B | ▲ | Wu *et al.* found that CDC25B was highly expressed in primary non-small cell lung cancer (NSCLC) | PMID: 9751615 |
| ADORA1 | - | No relevant reports were found | - |
| CHEK1 | - | No relevant reports were found. | - |
| NR3C1 | - | No relevant reports were found. | - |
| CDK2 | ▲ | Liu *et al.* demonstrated that CDK2 was highly expressed in LUAD tissues in comparison with normal tissues and high expression of CDK2 in LUAD has a poor prognosis. | PMID: 34409029 |
| ADORA3 | - | No relevant reports were found. | - |

**Table S22.** Literature Reports on Prognostic Genes of ARCR in COAD.

| **Gene** | **Feature** | **Summary** | **Citation** |
| --- | --- | --- | --- |
| PLK1 | ▲ | Raab *et al.* reported that in colon cancer, the overexpression of PLK1 is linked to more aggressive tumor behavior and a poorer prognosis. | PMID: 29549256 |
| ABL1 | - | No relevant reports were found. | - |
| NR3C1 | ▼ | Han *et al.* showed that increased miR-19b and decreased NR3C1 in colon cancer were correlated with poor prognosis and NR3C1 was directly targeted by miR-19b. | PMID: 34017210 |
| APP | ▲ | Venkataramani *et al.* showed that abundant APP staining was found in human pancreatic adenocarcinoma and colon cancer tissue | PMID: 20145244 |
| ESR1 | - | No relevant reports were found. | - |
| PIK3CA | - | No relevant reports were found. | - |
| VEGFA | ▲ | Esfahani *et al.* demonstrated that VEGFA were significantly upregulated in COAD tissues compare to normal tissues | PMID:38719941 |

**Table S23.** Literature Reports on Prognostic Genes of ARCR in BRCA.

| **Gene** | **Feature** | **Summary** | **Citation** |
| --- | --- | --- | --- |
| PLK1 | ▲ | Rizki *et al.* found that PLK1 was up-regulated in many invasive carcinomas and highly expressed in preinvasive *in sit*u carcinomas of the breast. | PMID: 18056432 |
| ADRA2C | ▼ | Sousa *et al.* found that ADRA2C was downregulated in BC tissues and correlated with metastatic BC and BC subtypes, thus the prognosis of the disease. | PMID: 36428611 |
| CDC25A | ▲ | Cangi *et al.* discovered that Cdc25A is overexpressed in primary breast tumors and this overexpression is associated with higher levels of cell cycle protein-dependent kinase 2 (Cdk2) enzyme activity in vivo | PMID: 10995786 |
| ADORA1 | ▲ | Mirza *et al.* showed that Adora1 is over-expressed in various breast cancer cell lines. | PMID: 16294023 |
| OPRD1 | ▲ | Montagnai *et al.* found that OGFR, OPRK1, and OPRD1 were upregulated in breast cancer tissue compared with normal tissue by bulk RNA-seq analysis of opioid receptors. | PMID: 33220939 |
| EGFR | ▲ | Masuda *et al.* demonstrated that overexpression of EGFR is observed in all subtypes of breast cancer | PMID: 23073759 |
| ESR1 | - | No relevant reports were found. | - |
| PIK3CA | - | No relevant reports were found. | - |
| HSP90AA1 | ▲ | Liu *et al.* demonstrated that the mRNA levels of HSP90AA1 in breast cancer tissues were higher than those in normal tissues. | PMID: 34109171 |
| AR | - | No relevant reports were found. | - |
| CDK1 | ▲ | Xing *et al.* demonstrated that CDK1, CCNA2, and CCNB1 gene expression levels were higher in BRCA compared with control tissue samples and were correlated with more-advanced tumor stage. | PMID: 33896262 |
| VEGFA | ▲ | Kawas *et al.* demonstrated that cytoplasmatic VEGFA/165b expression was higher in invasive breast cancer tumor cells than in normal tissues or stroma. | PMID: 35303290 |

**Table S24.** Literature Reports on Prognostic Genes of ARCR in STAD.

| Gene | Feature | Summary | Citation |
| --- | --- | --- | --- |
| ADRA2C | - | No relevant reports were found. | - |
| NR3C1 | - | No relevant reports were found. |  |
| EGFR | ▲ | Gao, *et al.* found that EGFR positive expression rate in the 78 cases of gastric cancer tissue was 57.7 % (45/78), while EGFR was not expressed in 20 cases of adjacent normal tissue, and the high EGFR expression was closely related to the incidence and development of gastric cancer. | PMID: 23608326 |
| ESR1 | - | No relevant reports were found. | - |
| AR | ▲ | Wang *et al.* demonstrated that AR expression was higher in Gastric Carcinogenesis tissues than in adjacent tissues.。 | PMID: 31413736 |
| CDK1 | ▲ | Zhang *et al.* demonstrated that the expression level of CDK1 in Gastric Carcinogenesis tissues was significantly higher than that in precancerous tissues and was significantly correlated with pathological stage and grade. | PMID: 33365314 |

**Table S25** Literature Reports on Prognostic Genes of ARCR in KIRC

| Gene | Feature | Summary | Citation |
| --- | --- | --- | --- |
| PLK | ▲ | Qian *et al.* indicated that PLK1 was validated to be highly expressed in ccRCC tissues and promoted ccRCC cell proliferation, migration, invasion, and cell cycle. | PMID: 35210814 |
| ADRA2C | - | No relevant reports were found. | - |
| SYK | - | No relevant reports were found. | - |
| APP | - | No relevant reports were found. | - |
| ESR1 | - | No relevant reports were found. | - |
| SRC | ▲ | Qayyum *et al.* demonstrated that Src was highly expressed in renal clear cell carcinoma. | PMID: 22814579 |
| PTGS2 | - | No relevant reports were found. | - |
| HSP90AA1 | - | No relevant reports were found. | - |
| AR | - | No relevant reports were found. | - |

**Table S26.** Literature Reports on Prognostic Genes of ARCR in ESCA.

| Gene | Feature | Summary | Citation |
| --- | --- | --- | --- |
| ADRA2B | - | No relevant reports were found. | - |
| CDC25A | ▲ | Li *et al.* demonstrated that CDC25A overexpression was implicated in promoting uncontrolled cell growth and division of esophageal cancer, particularly esophageal squamous cell carcinoma. | PMID: 37173384 |
| CDC25B | ▲ | Liu *et al.* demonstrated that high expression of CDC25B was detected in esophageal cancer cell lines and primary tumor tissues, but not in normal tissues. | PMID: 18722351 |
| CHEK1 | - | No relevant reports were found. | - |
| AKT1 | - | No relevant reports were found. | - |
| HSP90AA1 | - | No relevant reports were found. | - |

**Table S27** Literature Reports on Prognostic Genes of ARCR in THCA.

| Gene | Feature | Summary | Citation |
| --- | --- | --- | --- |
| PTPN1 | - | No relevant reports were found. | - |
| ADRA2C | - | No relevant reports were found. | - |
| CDC25A | ▲ | Ito *et al.* demonstrated that in thyroid cancer, CDC25A overexpression was associated with promoting cell cycle progression, cell proliferation, and tumor growth. | PMID: 12085185 |
| CHRM2 | - | No relevant reports were found. | - |
| CDC25B | ▲ | Ito *et al.* have demonstrated that CDC25B was frequently overexpressed in thyroid neoplasms, especially the follicular adenoma and minimally invasive follicular carcinoma. | PMID: 12085185 |
| APP | ▲ | Pietrzik *et al.* showed that the members of the APP family are highly expressed in follicle epithelial cells of the thyroid. | PMID： 9465092 |
| CDK2 | - | No relevant reports were found. | - |
| RAF1 | ▲ | Wang *et al.* showed that Raf1 was upregulated in thyroid cancer. | PMID: 26527888 |
| ADORA3 | - | No relevant reports were found. | - |
| ESR1 | ▲ | Yi *et al.* demonstrated that ESR1 gene expression and ESR ratio (ESR1/ESR2) were significantly higher in papillary thyroid cancer tissues than in paired normal thyroid tissues and caused a worse overall survival. | PMID:28124274. |
| PTGS2 | - | No relevant reports were found. | - |
| PIK3CA | - | No relevant reports were found. | PMID: 20804548 |
| HSP90AA1 | - | No relevant reports were found. | - |

▲: Gene was confirmed to be upregulated in this cancer leading to poor prognosis

▼: Gene was confirmed to be upregulated in this cancer leading to positive prognosis

**-:** No relevant reports were found regarding the relationship between the gene and this cancer

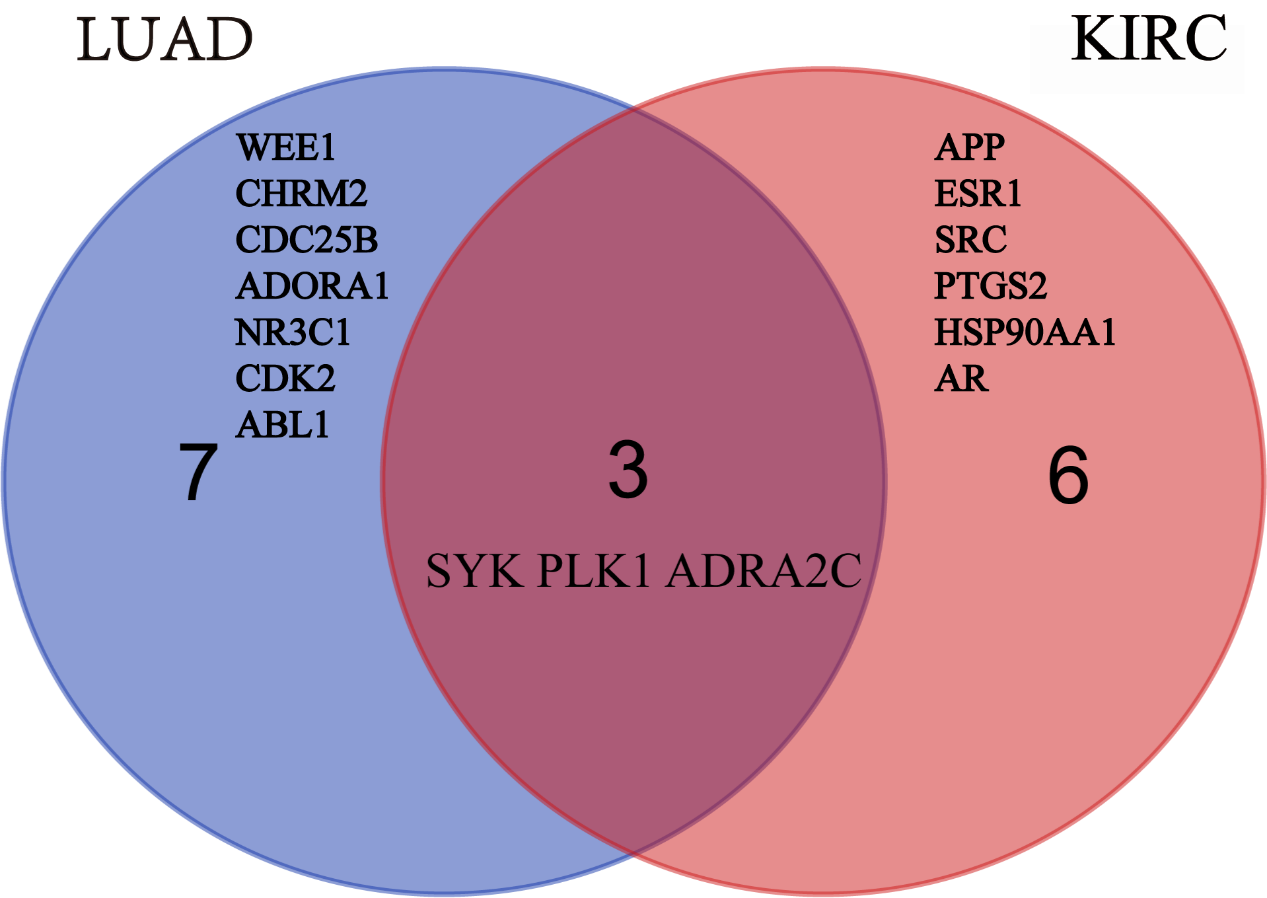

**Figure S1:** Intersection Genes in the Prognostic Gene Lists of LUAD and KIRC.

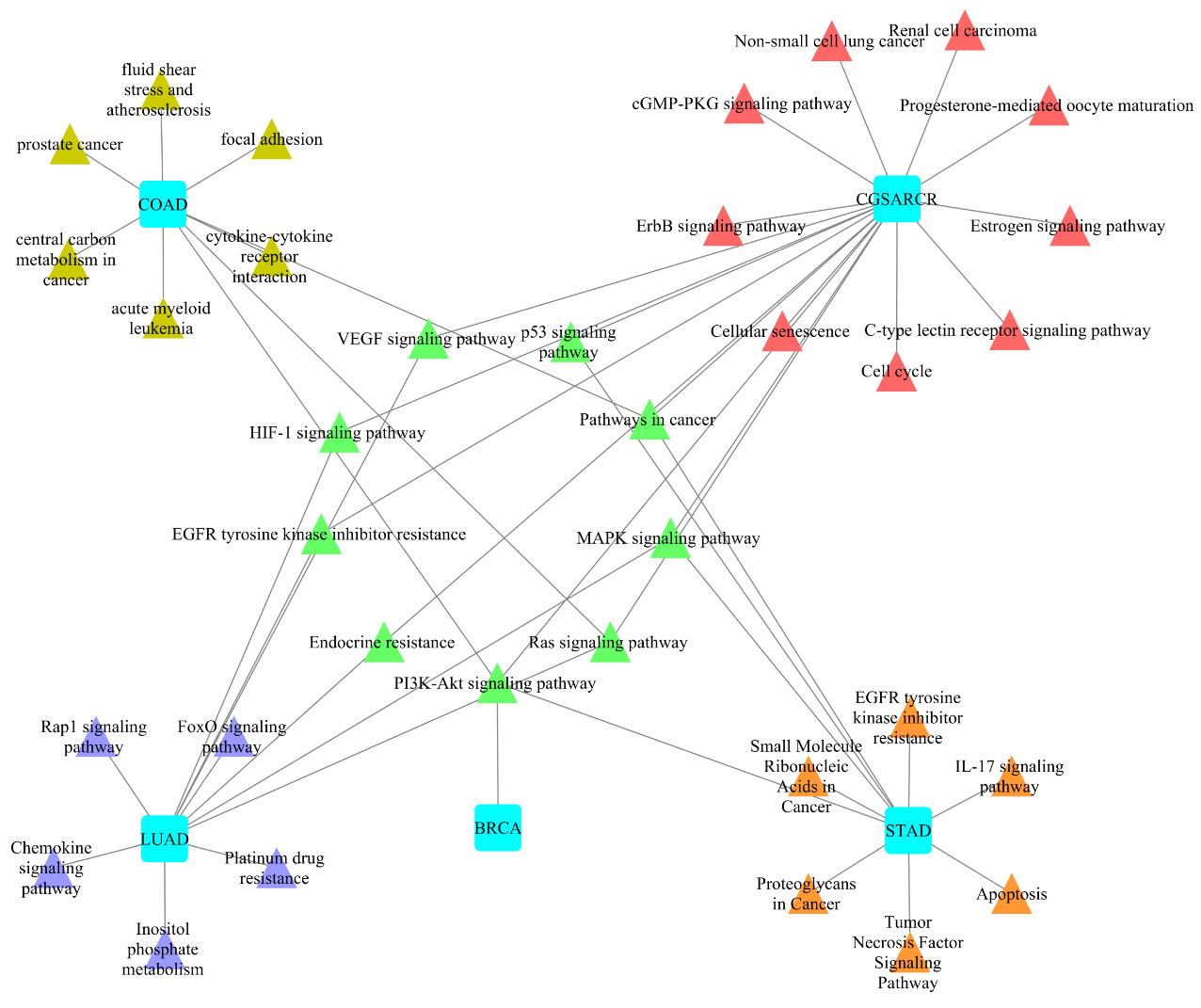

**Figure S2: Network of pathway similarities and differences between CGS_ARCR_ and the researched studies of LUAD, COAD, STAD, and BRCA.** The blue squares represent the lables of the studies for specific cancers and CGSARCR for our study, the green triangles indicate the same signaling pathways in the CGS_ARCR_ and the studies of four cancers, the red triangles are the signaling pathways independently owned by the CGS_ARCR_, the brown triangles are the signaling pathways independently owned by the ARCR in the study of COAD and the purple triangles are the signaling pathways independently owned by the ARCR in the study of LUAD, orange triangles are the signaling pathways independently owned by CGS_ARCR_ in the study of STAD. It is worth to note that ARCR in the BRCA with only one pathway experimentally verified.
